# Chemogenetic inhibition of IST1-CHMP1B interaction impairs endosomal recycling and promotes unconventional LC3 lipidation at stalled endosomes

**DOI:** 10.1101/2023.08.28.555152

**Authors:** Anastasia Knyazeva, Shuang Li, Dale P. Corkery, Kasturika Shankar, Laura K. Herzog, Xuepei Zhang, Birendra Singh, Georg Niggemeyer, David Grill, Jonathan D. Gilthorpe, Massimiliano Gaetani, Lars-Anders Carlson, Herbert Waldmann, Yao-Wen Wu

## Abstract

The Endosomal Sorting Complex Required for Transport (ESCRT) machinery constitutes a multisubunit protein complex that plays an essential role in membrane remodeling and trafficking. ESCRTs regulate a wide array of cellular processes, encompassing cytokinetic abscission, cargo sorting into multivesicular bodies (MVBs), membrane repair and autophagy. Given the versatile functionality of ESCRTs and the intricate organizational structure of the ESCRT complex, the targeted modulation of distinct ESCRT-mediated membrane deformations for functional dissection poses a considerable challenge. This study presents a pseudo-natural product targeting IST1-CHMP1B within the ESCRT-III complex. This compound specifically disrupts the interaction between IST1 and CHMP1B, thereby inhibiting the formation of IST1-CHMP1B copolymers essential for normal-topology membrane scission events. While the compound has no impact on cytokinesis, MVB sorting and exosome biogenesis, it rapidly hinders transferrin receptor (TfR) recycling in cells, resulting in the accumulation of transferrin in perinuclear endosomal recycling tubules. Stalled recycling endosomes acquire unconventional LC3 lipidation, establishing a link between non-canonical LC3 lipidation and endosomal recycling.

## Introduction

The Endosomal Sorting Complex Required for Transport (ESCRT) is an evolutionarily conserved protein machinery that plays a critical role in membrane remodeling events^1^ including multivesicular body (MVB) formation and exosome biogenesis^2–4^, autophagy^5–7^, viral budding^8^, cytokinesis^9^, nuclear envelope maintenance^1^^11^ and membrane repair^12–14^.

The ESCRT machinery consists of four complexes – ESCRT-0, -I, -II and -III with associated proteins, such as ALIX and VPS4. ESCRT-0, -I and -II play a central role in cargo sorting and in the recruitment and activation of ESCRT-III, which is directly involved in reshaping and severing membranes. In mammals, ESCRT-III comprises CHMP1-7 and IST1 proteins capable of forming polymers that drive membrane deformation. The AAA^+^ ATPase VPS4 binds to ESCRT-III subunits and disassembles ESCRT-III via ATP hydrolysis, generating the force for membrane scission ^15,16^. ESCRT-III proteins can bind to membranes to mediate both reverse- and normal-topology membrane remodeling processes^17^. While the majority of ESCRT-III proteins assemble into different structures on the membrane with a reverse topology, the IST1-CHMP1B complex can coat positively curved membranes to facilitate normal-topology membrane deformation^18–20^. IST1 plays an important role in endosomal tubulation^21,22^. However, IST1 and CHMP1B are also present at sites of reverse-topology membrane scission. The functioning of ESCRT-III complexes with distinct topologies in various membrane processes remains unclear. The sophisticate organization of the ESCRT complex and the multifaceted nature of its function collectively pose challenges in selectively modulating individual ESCRT-mediated membrane deformations for functional dissection.

Autophagy is an evolutionarily conserved catabolic process in eukaryotic cells. This process involves conjugation of ATG8 proteins to phospholipids on double-membrane structures known as autophagosomes, which then fuse with lysosomes to facilitate the degradation of cargo. Emerging evidence shows that the autophagy machinery also plays a role in several autophagy-independent processes. ATG8s can be conjugated to single-membrane compartments such as phagosomes and endolysosomes^23–28^. This non-canonical lipidation of ATG8s occurs independent of certain components of the core autophagy machinery. A prevalent mechanism of non-canonical ATG8 conjugation involves V-ATPase that directly binds ATG16L1 and recruits the ATG12-ATG5-ATG16L1 conjugation complex to membranes^29,30^. Non-canonical ATG8 conjugation has been shown to occur in response to different stimuli and its physiological roles have been demonstrated^31–34^. Further identification of non-canonical ATG8 conjugation under various stimuli will enhance understanding of its functions and mechanisms.

In this study, we identified IST1, a protein within the ESCRT-III complex, as the target of recently discovered pseudo-natural product Tantalosin. This compound induces non-canonical LC3 lipidation (LC3 being an ATG8 protein in mammals) in mammalian cells. Target engagement showed that the compound specifically disrupts the interaction between IST1 and CHMP1B, impeding the formation of the IST1-CHMP1B co-polymer. Remarkably, this compound does not affect the assembly of other ESCRT-III components. We found that the compound rapidly impairs ESCRT-dependent normal-topology membrane remodeling processes, leading to inhibition of TfR recycling without perturbating other ESCRT-medicated processes like cytokinesis, MVB sorting and exosome biogenesis. We showed that non-canonical LC3 lipidation occurs at stalled recycling endosomes in a V-ATPase dependent manner. Therefore, using a chemogenetic approach, we have dissected the functions of ESCRT-III with reverse and normal topologies. Additionally, we have identified a novel role of non-canonical LC3 lipidation in response to the inhibition of an ESCRT-III complex.

## Results

### Pseudo-natural product Tantalosin targets the ESCRT-III component IST1

We have conducted a phenotypic high-content screen (HCS) for small-molecule modulators of EGFP-LC3 lipidation, leading to identification of new regulators for autophagosome formation ^35,36^. Recently, we discovered macrocyclic Pseudo-Natural Products (PNPs) that induces LC3 lipidation in cells^37^. Tantalosin-I (hereafter referred as Tantalosin) was chosen for subsequent investigation (Figure 1A). To identify potential protein targets of Tantalosin, we performed the Proteome Integral Solubility Alteration (PISA) assay, a high-throughput approach based on thermal proteome profiling method ^38^. HEK293T cells were treated with 4 µM Tantalosin, 4 µM inactive compound (as control) or vehicle control (DMSO) for 1 hour. The inactive compound, chosen from the same compound class as Tantalosin, lacked the ability to induce LC3 lipidation (Supplementary Figure 1A). We analyzed proteome-wide shifts in solubility of samples treated with Tantalosin, comparing them to inactive compound or DMSO. In both instances, the ESCRT-III component IST1 emerged as a strong candidate that exhibited consistent and substantial engagement (Figure 1B). Importantly, no significant hits were identified in samples treated with the inactive compound in comparison to those treated with DMSO (Supplementary Figure 1B).

**Figure 1.**
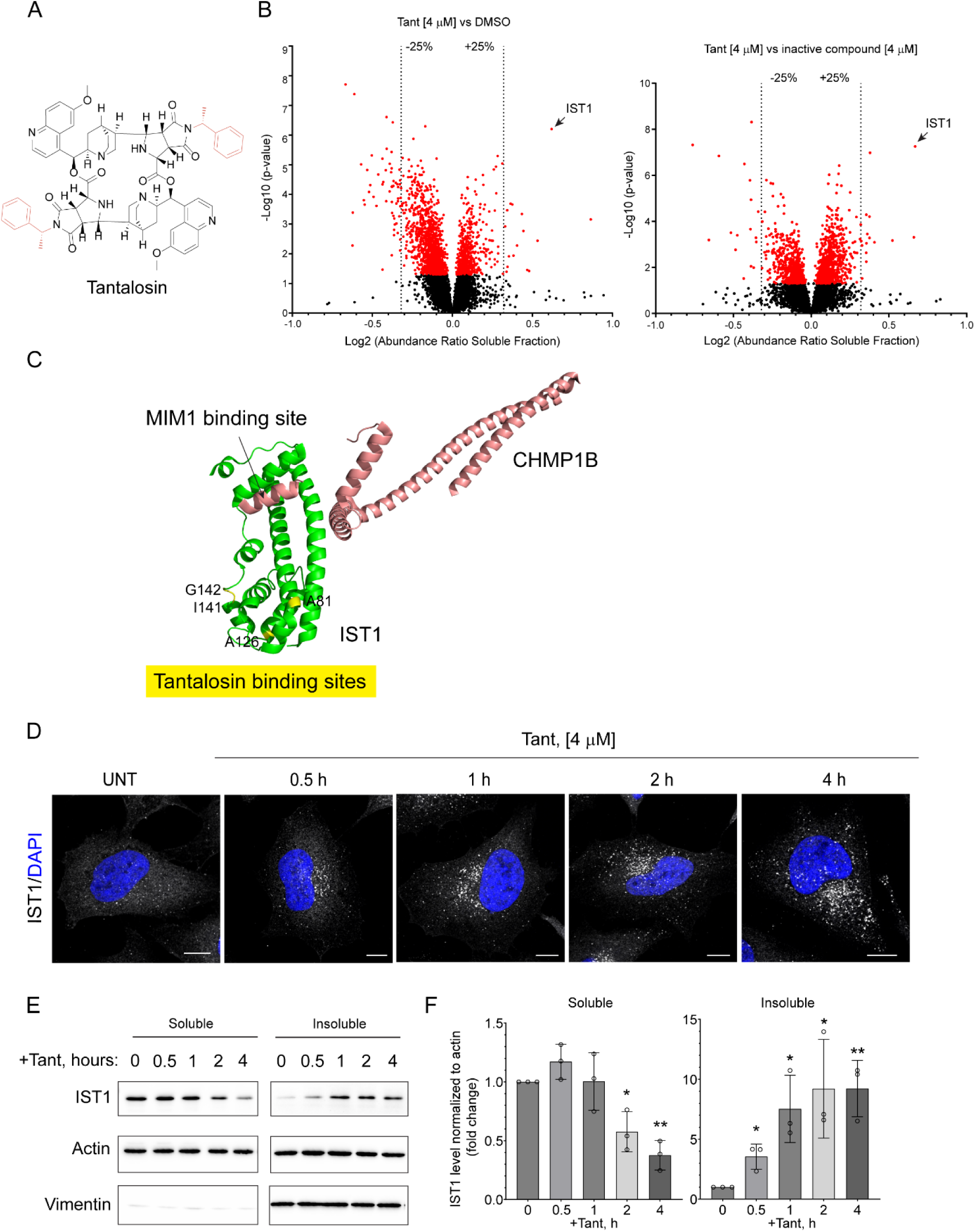
Identification of IST1 as a target of Tantalosin. **(A)** Chemical structure of Tantalosin. **(B)** Volcano plots of results obtained from the PISA assay performed in HEK 293T cells. Differences in solubility shifts are plotted against statistical significance. Results were shown as following: cells treated with 4 µM Tantalosin versus DMSO (left) and cells treated with 4 µM Tantalosin versus 4 µM inactive compound (right). **(C)** 3D structure of the IST1-CHMP1B complex within the co-polymer (PDB: 6TZ5^19^). IST1 (residues 1-189) and CHMP1B are shown in green and salmon red, respectively. Tantalosin-binding residues in IST1 are marked and highlighted in yellow. **(D)** HeLa cells were treated with 4 µM Tantalosin for indicated time and immunostained for endogenous IST1. Nuclei were counterstained with DAPI. Scale bar: 10 µm. Results are representative of three independent experiments. **(E)** Western blot analysis of IST1 protein level in NP40-soluble and NP40-insoluble fraction of protein lysates prepared from HeLa cells treated with 4 µM Tantalosin for indicated time. Vimentin serves as a control for the insoluble fraction. **(F)** Quantification of IST1 levels from (E) normalized to actin level. Fold change is quantified relative to time point “0 h” for each of soluble and insoluble fractions. All data presented as graph bars with mean ± SD for three independent experiments, individual values for each experiment are presented as dots. * P < 0.05, ** P < 0.005.

To further validate the targeting of IST1 by Tantalosin, we carried out hydrogen-deuterium exchange mass spectrometry (HDX-MS). This technique enabled us to explore potential binding sites of Tantalosin within IST1. We identified the minimal sequence segment displaying differential uptake, which covered all peptides spanning the protein sequence in IST1 treated with both Tantalosin and DMSO at varying time intervals. Tantalosin led to an alteration in HDX of amino acid residues 79-89 and 136-146 (Figure 1C; Supplementary Figure 2A). In the case of residues 79-89, a time-dependent decrease in differential deuterium uptake occurred at residue A81. This reduction suggests its involvement in the binding to Tantalosin, which also appeared to cause the exposure of the adjacent residue F83 (Supplementary Figure 2B). Similarly, for residues 136-146, a dynamic pattern of decreased differential deuterium uptake was evident, peaking at 150 seconds on residues I141 and G142. These residues are adjacent to the residue A126, which also exhibited a similar pattern. Upon binding to Tantalosin, the neighboring residue R147 became exposed (Supplementary Figure 2A). Since these engaged residues do not reside within a single region (Figure 1C), the interaction between Tantalosin and IST1 might induce a conformational change in IST1, potentially generating a new binding site for Tantalosin.

To investigate the impact of Tantalosin on IST1 in cells, we subjected HeLa cells to a treatment with 4 µM Tantalosin. Tantalosin induced rapid accumulation of endogenous IST1 into punctate structures within the cell in a time-dependent manner (Figure 1D). Conversely, no accumulation of IST1 was observed in cells treated with the inactive compound (Supplementary Figure 1C). Cell lysate membrane fractionation revealed a decrease in IST1 levels in the detergent-soluble fraction and an elevation in the detergent-insoluble fraction upon Tantalosin treatment (Figure 1E, F). Collectively, these data suggested that IST1 is the cellular target of Tantalosin.

### Tantalosin specifically impairs IST1-CHMP1B interaction and copolymer formation

IST1 interacts with CHMP1B in solution with a low affinity^20,39^. Using micro-scale thermophoresis (MST), we determined that recombinant full-length IST1 binds to CHMP1B with a dissociation constant (K_D_) of 9.0 ± 2.4 µM (Figure 2A). We observed a dose-dependent inhibition of the IST1-CHMP1B interaction by Tantalosin. This is evident from the increase in K_D_ from 20.3 ± 0.8 µM (with 1.8 µM Tantalosin) to 153.3 ± 1.0 µM (with 3.5 µM Tantalosin) and complete inhibition of binding with 7 µM Tantalosin, indicating that Tantalosin effectively blocks the interaction of IST1 and CHMP1B in solution. Employing a non-competitive inhibition model, we fitted the IST1-CHMP1B binding data in the presence and absence of Tantalosin, resulting in a Tantalosin inhibition constant (K_I_) of 3.5 µM (Supplementary Figure 3A). Conversely, we did not observe any inhibitory effect on the IST1-CHMP1B interaction upon treatment with the inactive compound (Supplementary Figure 3B).

**Figure 2.**
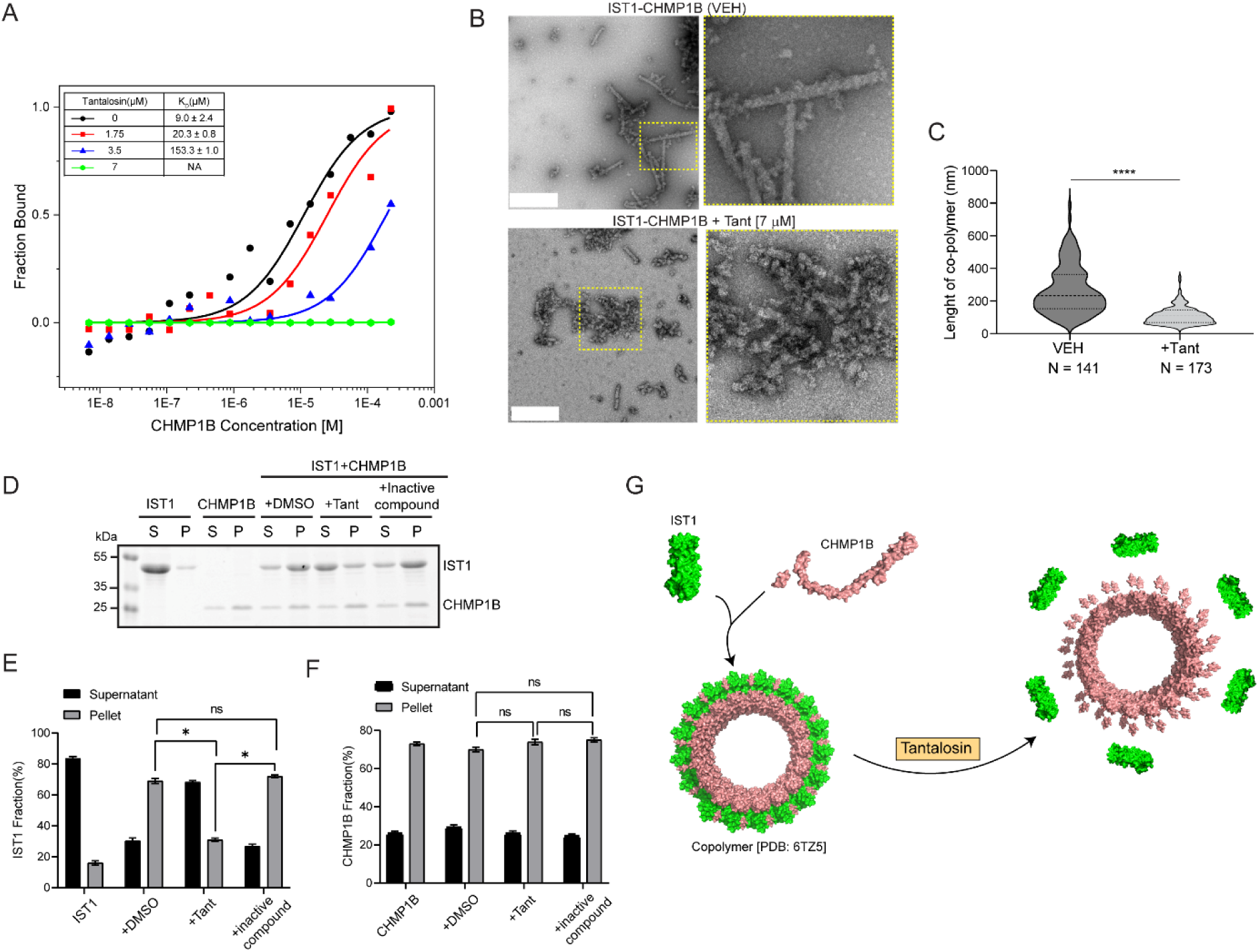
Tantalosin inhibits the interaction and assembly of IST1 and CHMP1B. **(A)** Dose-dependent binding curves of IST1 and CHMP1B in solution in the presence of indicated concentrations of Tantalosin. CHMP1B (225 µM to 6.8 nM) was mixed with IST1 (80 nM) in assay buffer. Inset represents a table of mean K_D_ ± SD for the IST1-CHMP1B binding with 0-7 µM Tantalosin from three independent experiments; NA: not available. **(B)** Representative TEM images of the IST1-CHMP1B co-polymer treated with vehicle (DMSO) or 7 µM Tantalosin. Scale bar: 0.5 µm. **(C)** Quantification of the length of IST1-CHMP1B co-polymer treated with vehicle (DMSO) or 7µM Tantalosin. N represents the total number of analyzed IST1-CHMP1B co-polymers from three independent experiments; **** P <0.0001. **(D)** SDS-PAGE analysis of IST1 (5 µM) and CHMP1B (5 µM) co-polymer sedimentation assay with DMSO, 7 µM Tantalosin or 7 µM inactive compound. S: supernatant; P: pellet. **(E)** and **(F)** Quantification of IST1 and CHMP1B levels, respectively, shown in (D) from three independent experiments; *P <0.05, ns: not significant. **(G)** Demonstration of the inhibition of the IST1-CHMP1B co-polymer (PDB: 6TZ5^19^) by Tantalosin. IST1 and CHMP1B are shown in green and salmon red, respectively.

HDX-MS revealed the involvement of I141 and G142 of IST1 in Tantalosin binding (Supplementary Figure 2A). To validate this finding, I141 or G142 was mutated to aspartic acid, respectively. In both cases, the IST1 mutants exhibited comparable binding affinities to CHMP1B, regardless the presence or absence of Tantalosin (Supplementary Figure 3C, D). Therefore, the mutation of I141 or G142 disrupted the Tantalosin binding, thereby abolishing its inhibitory effect on the IST1-CHMP1B interaction. This result confirmed the association of Tantalosin with I141 and G142 of IST1. Moreover, Tantalosin did not engage in competition with CHMP1B for binding at these residues, in keeping with the IST1-CHMP1B structure ^18–20^. Instead, the binding of Tantalosin to I141 and G142 presumably induces a conformational change in IST1, preventing the interaction with CHMP1B (Figure 1C).

IST1 and CHMP1B can assemble into helical tubes, and current models propose that IST1 decorates the outer layer of the CHMP1B tube ^18–20^. Having verified Tantalosin’s inhibitory effect on the IST1-CHMP1B interaction in solution, we proceeded to examine the impact of Tantalosin on the assembly of the IST1-CHMP1B co-polymer. Examination by transmission electron microscopy (TEM) revealed that Tantalosin disrupts the formation of ordered IST1-CHMP1B co-polymers (Figure 2B), leading to a significant reduction in their length compared to the control (Figure 2 B, C). To further validate inhibition of IST1-CHMP1B co-polymer formation, we executed a sedimentation assay. The assay showed that the insoluble fraction (co-polymers) of IST1 decreases with the addition of Tantalosin, while remaining unaffected by the inactive compound (Figure 2D, E). Notably, the distribution of CHMP1B in the soluble and insoluble fractions remained unaffected by Tantalosin (Figure 2D, F). These findings suggest that Tantalosin inhibits the formation of IST1-CHMP1B co-polymers through its interference with the binding between IST1 and CHMP1B (Figure 2G).

The IST1-CHMP1B complex adopts a normal topology, coating on positively curved membranes ^18–20^. Conversely, other ESCRT-III proteins like CHMP2A and CHMP3 can co-polymerize to form helical tubes that mediate reverse-topology membrane deformation^40^. Notably, TEM analysis and sedimentation assays demonstrated that Tantalosin does not influence the formation of CHMP2A-CHMP3 co-polymers (Supplementary Figure 4). Hence, Tantalosin specifically hampers the assembly of the IST1-CHMP1B complex – a member of the ESCRT-III complex that embraces a normal topology on membranes.

The C-terminal MIT-interacting motif (MIM) 1 of CHMP1B is involved in its interaction with IST1^20^. Interestingly, the presence of Tantalosin did not lead to a significant inhibition in the binding between the CHMP1B MIM1 peptide (Supplementary Figure 3E). Additionally, IST1 features both a C-terminal MIM1 and MIM2 (residues 321-366), which associate with the MIT domains of ESCRT proteins (VPS4, LIP5 and MITD1) as well as spastin^41–43^. Tantalosin did not affect the binding between IST1 and VPS4A (Supplementary Figure 3F), in line with the HDX-MS mapping of the Tantalosin-binding regions on IST1 (Figure 1C). Therefore, Tantalosin employs a unique interaction mode with IST1 to specifically disrupt the IST1-CHMP1B interaction. This property could establish Tantalosin as a mechanism-specific chemogenetic tool for inhibiting the IST1-CHMP1B complex-mediated normal-topology membrane remodeling.

### Tantalosin does not affect cytokinesis, MVB sorting and exosome biogenesis

IST1 and CHMP1B are present at the midbody and play an essential role in cytokinesis^41,42,44^. It is not clear how the normal-topology structure of the IST1-CHMP1B co-polymer could participate in cytokinesis ^17^. Consistent with previous reports^41,42^, silencing IST1 using siRNA in HeLa cells led to a significant increase in the count of cells connected by midbodies, indicating compromised cytokinesis (Supplementary Figure 5A-C). However, Tantalosin did not impair cytokinesis (Supplementary Figure 5A-C), aligning with our finding that Tantalosin does not interfere with the interaction between IST1 and VPS4A (Supplementary Figure 3D). These results suggest that the selective inhibition of the IST1-CHMP1B co-polymer, achieved through Tantalosin treatment, does not prevent IST1 from associating with other ESCRT-III proteins (e.g. VPS4A) that are required for the reverse-topology membrane scission during cytokinesis. Therefore, cytokinesis does not require the IST1-CHMP1B complex-mediated normal-topology membrane remodeling.

Next, we examined the effect of Tantalosin on multivesicular body (MVB) sorting and exosome biogenesis. Following endocytosis, Epidermal Growth Factor Receptor (EGFR) is sorted into the intraluminal vesicles of MVBs via an ESCRT-dependent membrane scission event^45,46^. Subsequently, MVBs fuse with lysosomes to facilitate EGFR degradation. Tantalosin treatment had no impact on EGFR degradation upon EGF stimulation (Supplementary Figure 5D, E), indicating that receptor sorting into MVBs is not impaired. Alternatively, MVBs can fuse with the plasma membrane, leading to the release of intraluminal vesicles (exosomes) into the extracellular space^47^. Purified extracellular vesicles (EVs) from conditioned media of Tantalosin-treated cells exhibited unchanged expression of exosome markers (ALIX and TSG101) (Supplementary Figure 5F) and EVs morphology (Supplementary Figure 5G). Therefore, Tantalosin-induced inhibition of the IST1-CHMP1B co-polymer does not affect MVB sorting mediated by reverse-topology membrane deformation.

### Tantalosin inhibits endosomal recycling

Knockdown of IST1 causes impaired TfR recycling, implicating IST1 and its binding partner spastin in endocytic recycling regulation ^21,48^. However, it remains unclear whether IST1’s role in endocytic recycling is governed by the formation of IST1-CHMP1 co-polymers, its engagement with other ESCRT proteins and spastin, or a combination of both. Given that Tantalosin specifically impedes IST1-CHMP1 co-polymer without affecting IST1’s interaction with the MIT-containing protein VPS4A, Tantalosin could serve as a chemogenetic tool to address this question. Treatment of HeLa cells with Tantalosin led to a time-dependent accumulation of endogenous TfR, accompanied by tubular structure formation (Figure 3A). Notably, IST1-positive vesicles co-localized with TfR containing tubules, indicating IST1’s recruitment to the tubules persists, yet the complex was incapable of performing scission (Figure 3A, B). This observation was confirmed by treating cells with fluorescently labeled transferrin (Tf-555), showing similar accumulation in tubular structures following Tantalosin treatment (Figure 3C). This recycling defect extends beyond TfR, as EGFR similarly accumulated in the perinuclear region (Supplementary Figure 6).

**Figure 3.**
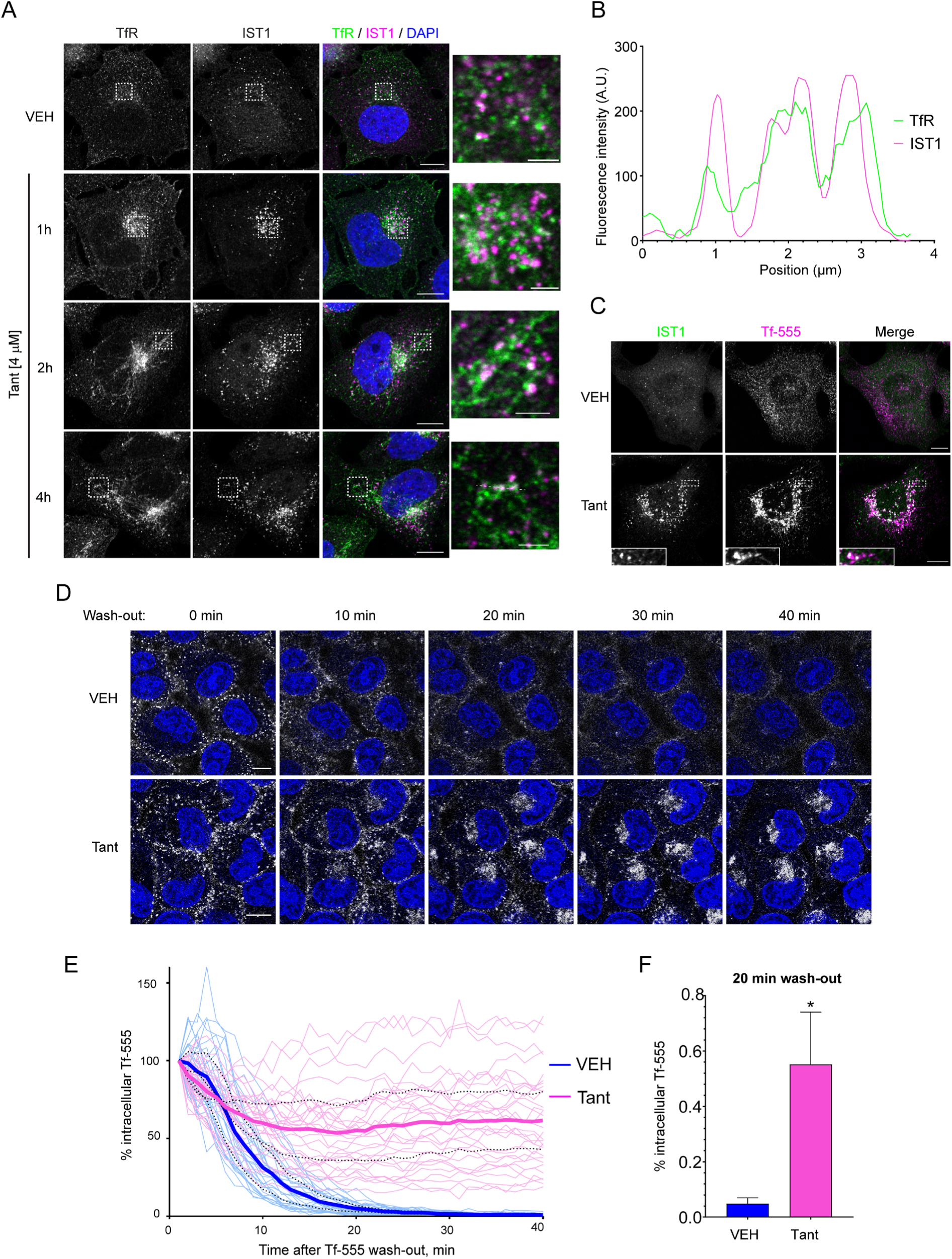
Tantalosin inhibits endosomal recycling. **(A)** Representative confocal images of HeLa cells treated with 4 µM Tantalosin for indicated time and immunostained for endogenous transferrin receptor (TfR) and IST1. Insets represent regions with transferrin-positive tubules. Scale bars: 10 µm for whole images and 2 µM for insets. **(B)** Fluorescence intensity profiles of TfR (green) and IST1 (magenta) for inset region of 4 h Tantalosin treatment marked by dashed line. (**C**) Representative images of HeLa cells treated with transferrin conjugated to Alexa Fluor 555 (Tf-555), treated with 4 µM Tantalosin for 2 h and immunostained for IST1. Scale bars: 10 µM. **(D)** Representative live-cell confocal images of HeLa cells pre-treated with VEH or 4µM Tant, pre-labeled with transferrin conjugated with Alexa Fluor 555 (Tf-555), and then chased with Tf-555-free media (wash-out) for indicated time. Nuclei were labeled with DRAQ5. Scale bars: 10 µm. **(E)** Quantification of percentage of remaining intracellular Tf-555 area per cell, normalized to total cell area relative to time point “0 min” (set as 100%) for images acquired in (D). Light lines represent individual cells. Bold lines represent mean value for three independent experiments, dashed lines represent SD. **(F)** Quantification of percentage of remaining Tf-555 after 20 min of Tf-555 wash-out for experiment described in (D). Bar graph represents mean ± SD for three independent experiments. Significance was determined using two-tailed Student t-test. * P < 0.05.

We explored the disruption of TfR recycling through a transferrin pulse-chase assay, where HeLa cells were pretreated with fluorescently labeled transferrin, then subjected to wash-out in the presence or absence of Tantalosin. Vehicle treated cells displayed a near-complete disappearance of the Tf-555 signal within 30 minutes of wash-out, attributed to recycling and subsequent detachment of fluorescently labelled ligand. Conversely, Tantalosin treatment caused rapid accumulation of labelled transferrin in the perinuclear region, a scenario that persisted throughout the experiment (Figure 3D-F). These findings suggest that Tantalosin acts as a potent chemical inhibitor of endosomal recycling. Hence, the normal-topology IST1-CHMP1 co-polymer is required for endosomal recycling, in line with its function of membrane tubulation, constriction and thinning ^19,49^.

### Tantalosin induces LC3 lipidation on stalled recycling endosomes

In a previous study, we screened a family of 20-membered macrocyclic PNPs in an autophagy induction assay, pinpointing Tantalosin as the most robust LC3 lipidation inducer ^37^. To explore a potential correlation between inhibition of endocytic recycling and LC3 lipidation, we co-treated MCF7 cells stably expressing EGFP-LC3 with fluorescently labelled transferrin and Tantalosin. Interestingly, LC3 accumulated at endocytic recycling compartments marked with Tf-555 after Tantalosin treatment (Figure 4A-B). This observation was further validated by co-staining for endogenous LC3 and TfR in

**Figure 4.**
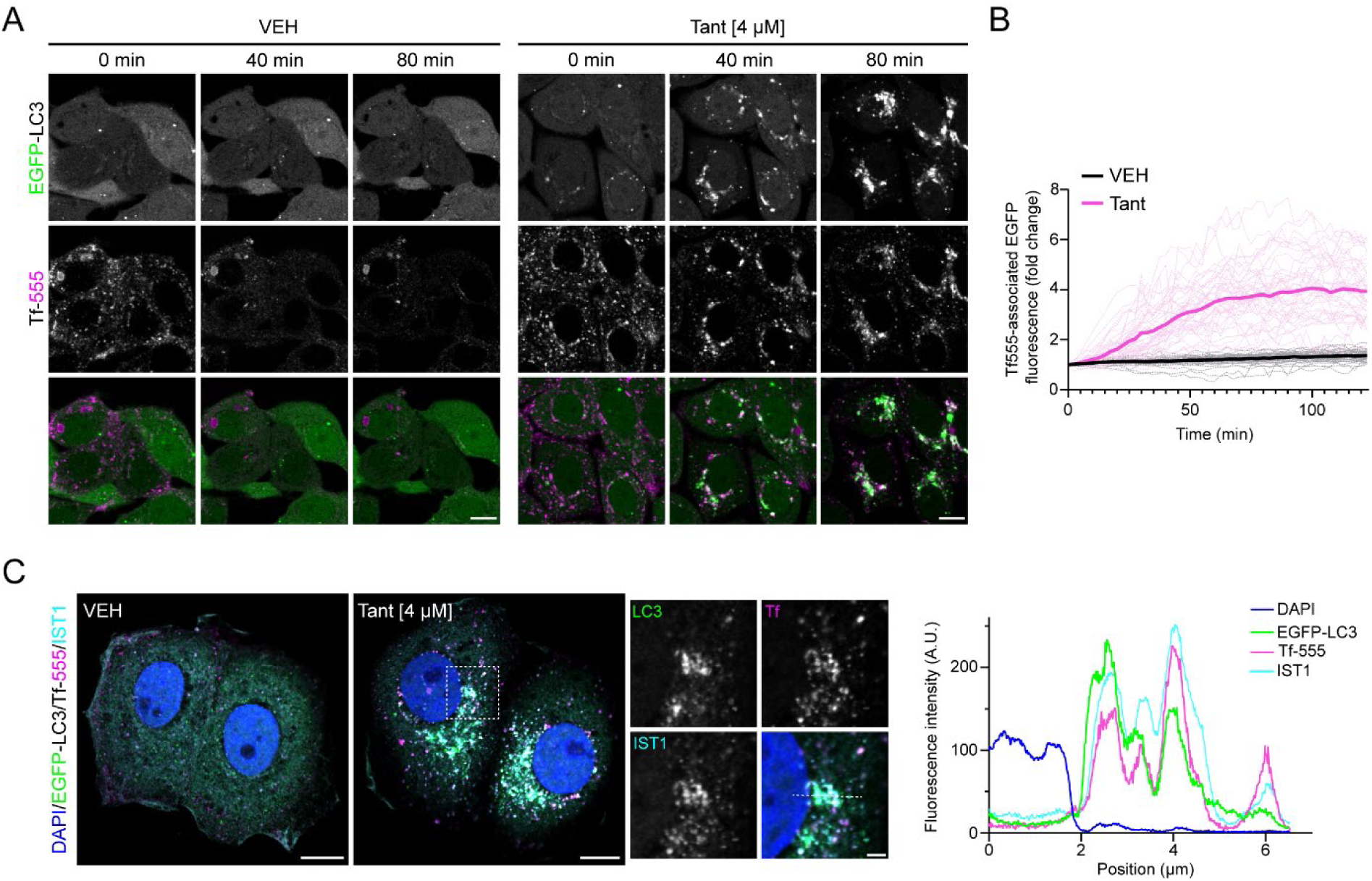
Tantalosin induces LC3 lipidation associated with transferrin receptor. **(A)** Representative live-cell confocal images of MCF7 cells stably expressing EGFP-LC3 (green), pre-labeled with fluorescent transferrin (Tf-555, magenta), and pre-treated with VEH or 4 µM Tant for 30 min prior to the start of imaging. Live-cell imaging was acquired in presence of VEH or 4 µM Tant respectively. Time indicates the duration of imaging. Scale bar: 10 µm. **(B)** Quantification of Tf-555-assosiated EGFP-LC3 fluorescence intensity per cell normalized to time point “0 min” for VEH- or Tantalosin-treated cells respectively. Light lines represent individual cells pooled from three independent experiments. Bold lines represent mean values of three independent experiments. **(C)** Representative confocal images of MCF7 cells stably expressing EGFP-LC3 treated with Tf-555 and 4 µM Tant for 2 h and stained for endogenous IST1 (left). Corresponding fluorescence intensities profiles for each channel are presented (right). Scale bars: 10 µm for original images and 2 µm for insets.

Tantalosin-treated HeLa cells (Supplementary Figure 7A). Immunostaining for IST1 confirmed the co-localization of Tf-555, LC3 and IST1 within the same compartment following Tantalosin-mediated recycling inhibition (Figure 4C). This indicates that impaired IST1 function on tubular recycling endosomes may promote the conjugation of LC3 onto stalled endosomal membranes. Intriguingly, IST1 depletion, previously shown to enhance endosomal tubulation^21^, did not lead to LC3 accumulation. This implies that genetic knockdown might promote compensatory mechanisms and/or assembly of CHMP1B-binding deficient IST1 on recycling tubules acts as a prerequisite for LC3 lipidation (Supplementary Figure 7B, C).

To further examine the causality between LC3 lipidation and endocytic recycling defects, we used LC3 lipidation-deficient ATG7 knock-out (KO) cells. Robust accumulation of transferrin was observed in ATG7 KO cells treated with Tantalosin (Supplementary Figure 8), suggesting that LC3 lipidation does not promote the endocytic recycling defect observed during Tantalosin treatment. Altogether, these data confirm that Tantalosin induces LC3 lipidation on stalled recycling endosomes, a process associated with aberrant accumulation of IST1 on membranes.

### Tantalosin induces V-ATPase-dependent non-canonical LC3 lipidation

LC3 lipidation is considered a hallmark of canonical autophagy - a lysosome-dependent catabolic pathway used for intracellular cargo degradation. In our previous study, we observed that inhibition of autophagosome-lysosome fusion did not result in an additive accumulation of lipidated LC3 in Tantalosin-treated cells^37^. This indicates that the Tantalosin-induced lipidated LC3 could be involved in a lysosome-independent, non-degradative function. Supporting this hypothesis, we found that TfR protein levels remain unaltered after Tantalosin treatment, and co-treatment with the autophagosome-lysosome fusion inhibitor, chloroquine, does not lead to elevated TfR levels (Figure 5A, B). Therefore, the LC3 lipidation induced by Tantalosin is not a consequence of canonical autophagy induction intended for TfR degradation.

**Figure 5.**
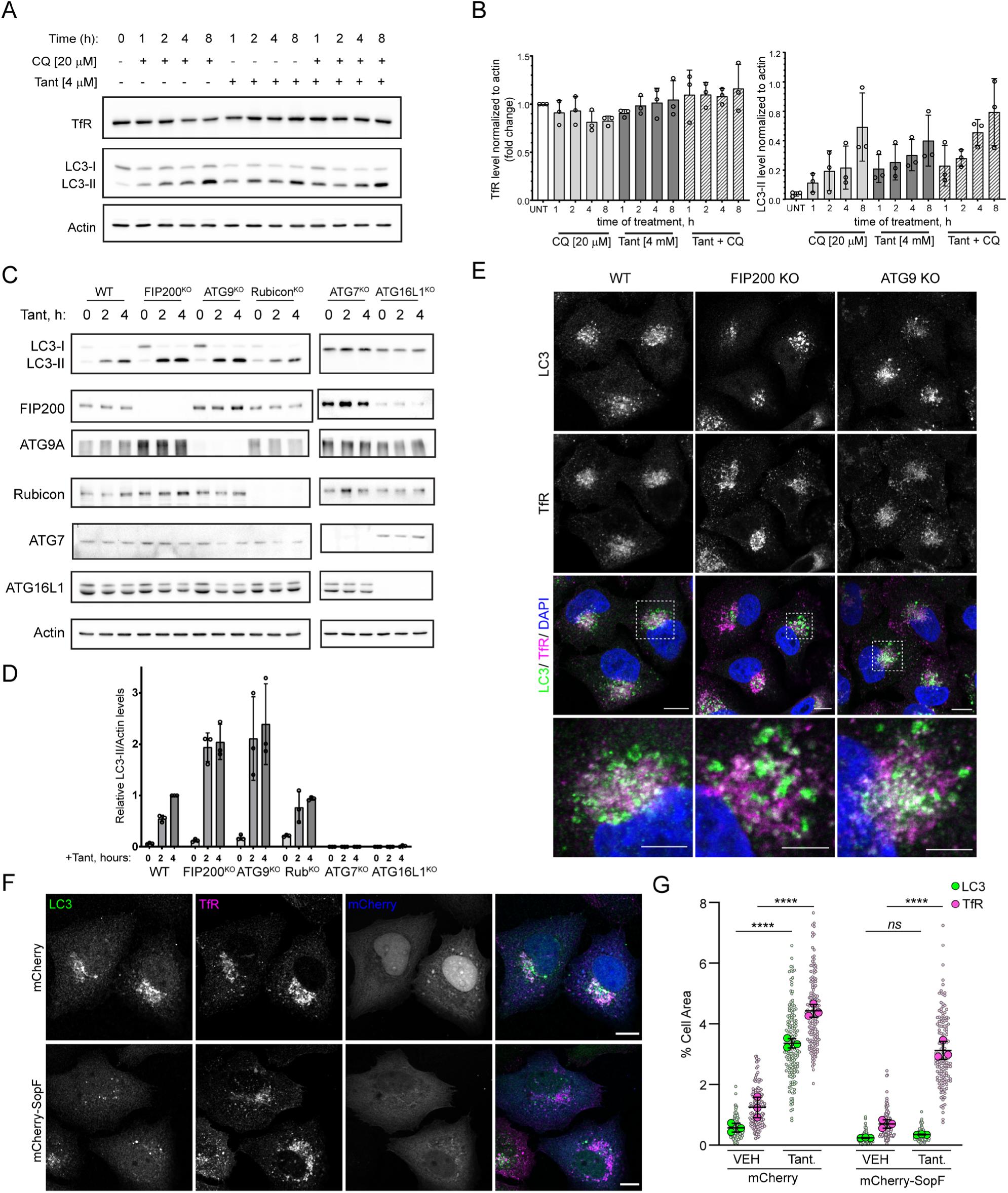
Tantalosin induces non-canonical LC3 lipidation. **(A)** Western blot analysis of transferrin receptor (TfR) and LC3 levels in cells treated with 4 µM Tantalosin and/or 20 µM chloroquine (CQ) for indicated time. **(B)** Quantification of TfR (upper panel) and LC3-II (lower panel) levels obtained from Western blot experiments presented in (A). TfR levels were normalized to actin levels, and fold-change was quantified relatively to untreated (UNT) samples. LC3-II level was normalized to actin. Bar graph represents mean ± SD for three independent experiments. **(C)** Western blot analysis of LC3 lipidation status in HeLa WT, FIP200 KO, ATG9 KO, Rubicon KO, ATG7 KO and ATG16L1 KO cell lines treated with 4 µM Tantalosin for indicated time. Verification of knock-out for each cell line is presented. **(D)** Quantification of LC3-II levels from experiments presented in (C). Data is normalized to actin and fold change is quantified relative to LC3-II levels in WT cells treated with Tantalosin for 4 h. Bar graph represents mean ± SD for three independent experiments. **(E)** Representative confocal images of WT, FIP200 KO and ATG9 KO HeLa cells treated with 4 µM Tantalosin for 4 h and immunostained for LC3 and transferrin receptor (TfR). Nuclei were counterstained with DAPI. Scale bars: 10 µm for original images and 2 µm for insets. **(F)** Representative images of FIP200 KO HeLa cells transiently transfected with mCherry or mCherry-SopF for 24 hs, treated with 4 µM Tantalosin for 4 h and immunostained for LC3 and TfR. Scale bars: 10 µm. (**G**) Quantification of TfR and LC3 cell area from (F). Small points represent individual cells from three independent experiments. Large points represent the mean of individual experiments (n = 50 cells per experiment). Bars represent the mean ± SD from the three experiments. Significance was determined from biological replicates using one-way ANOVA with Tukey’s multiple comparisons tests. ****P < 0.0001, ns: not significant.

To characterize the nature of LC3 lipidation induced by Tantalosin, we treated a panel of HeLa cell lines deficient for a variety of autophagy-related genes (FIP200, ATG9, Rubicon, ATG7 and ATG16L1) with Tantalosin and assessed LC3 lipidation via Western blot. Cells deficient for proteins involved in autophagosome initiation (FIP200 KO and ATG9 KO) exhibited lipidated LC3 levels comparable to those in wild type (WT) cells (Figure 5C, D), suggesting that LC3 is not conjugated to canonical autophagosomes. Similarly, LC3 lipidation remained unaffected in Rubicon KO cells, suggesting that Tantalosin-induced LC3 lipidation is distinct from LC3-assosiated phagocytosis and endocytosis – processes dependent on Rubicon^50^. However, ATG7 KO and ATG16L1 KO cells treated with Tantalosin exhibited significant inhibition of LC3 lipidation, indicating that Tantalosin-induced LC3 lipidation is dependent on the ATG conjugation systems (Figure 5 C, D). Notably, canonical autophagy-deficient cells lines (FIP200 KO and ATG9 KO) exhibited identical LC3 distribution patterns, co-localizing with unrecycled transferrin receptor after Tantalosin treatment (Figure 5E, Supplementary Figure 9). Collectively, these findings reveal a novel non-canonical form of LC3 lipidation linked to stalled recycling endosomes as a result of impaired IST1-CHMP1B interaction.

Non-canonical ATG8 conjugation has been shown to be dependent on the vacuolar-type H^+^-ATPase (V-ATPase) which directly binds ATG16L1 and recruits the ATG12-ATG5-ATG16L1 conjugation complex to target membranes^29,30,51^. The *Salmonella* T3SS effector SopF, has been demonstrated to hinder ATG16L1 recruitment to V-ATPase, thereby impeding non-canonical LC3 lipidation without inhibiting endosomal acidification^52^. Interestingly, transient expression of SopF inhibited Tantalosin-induced LC3 lipidation, as assessed in FIP200 KO cells lacking canonical autophagy (Figure 5F, G). Taken together, Tantalosin induces V-ATPase-dependent non-canonical LC3 lipidation on stalled recycling endosomes.

## Discussion

This study describes the identification and characterization of a novel PNP targeting the IST1-CHMP1B interaction within the ESCRT-III complex, thereby specifically disrupting endosomal recycling. The binding sites of Tantalosin on IST1 are located outside the IST1-CHMP1B interface, suggesting that Tantalosin acts as an allosteric inhibitor of the IST1-CHMP1 interaction. Tantalosin may induce a conformational change of IST1, resulting in disruption of the IST1-CHMP1 co-polymer. Tantalosin impairs neither the interaction of CHMP1B MIM1 with IST1 nor the interaction between IST1 and MIT-containing proteins like VPS4. Furthermore, Tantalosin does not interfere with the assembly of ESCRT-III complexes involved in reverse-topology membrane scission processes, such as the CHMP2-CHMP3 complex. Therefore, Tantalosin specifically inhibits the formation of the IST1-CHMP1B complex on membranes, while preserving the interaction between IST1 and other ESCRT components. Consequently, our findings establish Tantalosin as a novel chemogenetic tool for investigating the function of the ESCRT-III complex in both normal- and reverse-topology membrane scission events.

Perturbation of the ESCRT-III complex has been achieved by genetic depletion of its components or through overexpression of mutant VPS4 (E228Q)^53^. Due to the sophisticated organization of the ESCRT complex, genetic perturbation of individual components could potentially interfere with complex assembly, leading to global inhibition of ESCRT-dependent processes. For example, knockdown of IST1 not only disrupts the IST1-CHMP1B complex but also hampers the recruitment of VPS4, MITD1 and spastin. IST1 knockdown simultaneously causes defects in cytokinesis, endosomal tubulation and autophagy^7,21,41,42^. Tantalosin enables us to rapidly and precisely inhibit the IST1-CHMP1B complex. We demonstrate that the normal-topology membrane scission medicated by the IST1-CHMP1 complex is dispensable for cytokinesis, MVB sorting and exosome biogenesis. These processes are probably driven by a reverse-topology membrane scission. The role of IST1 in cytokinesis may be largely attributed to the recruitment and activation of other ESCRT components such as AAA^+^ ATPase VPS4 *via* its MIM motif ^41,43^.

On the other hand, we show that the IST1-CHMP1 complex-mediated normal-topology membrane scission plays an essential role in endosomal recycling. The IST1-CHMP1 co-polymer drives membrane tubulation, constriction and thinning ^19,49^. The IST1-CHMP1 co-polymer may coat endosomal tubes and collaborates with another AAA^+^ ATPase spastin to facilitate membrane scission ^21,22^. Therefore, ESCRT proteins execute normal- and reverse-topology membrane scissions, each playing distinct roles in different membrane processes. Investigating the effect of Tantalosin on other membrane processes, such as lipid droplet transportation to peroxisomes or peroxisome scission from the endoplasmic reticulum will provide insights into the role of the IST1-CHMP1B complex in these processes ^54,55^.

Tantalosin’s specificity in inhibiting the IST1-CHMP1B complex and disrupting endosomal recycling highlights its potential to target receptors and adhesion molecules essential for cancer development and progression ^56^. Previously, an endosomal recycling inhibitor primaquine was shown to interfere with EGFR endosomal trafficking and effectively inhibit proliferation of breast cancer cells ^57–59^. Expanding the repertoire of chemical modulators targeting the endocytic pathway with distinct mechanisms of action can be beneficial for the development of novel anti-cancer therapeutics.

Tantalosin was initially identified as an inducer of LC3 lipidation. We expand upon this finding to show that Tantalosin induces non-canonical LC3 lipidation on stalled recycling endosomes. Tantalosin-induced LC3 lipidation is dependent on the ATG conjugation systems but occurs independently of FIP200 and ATG9A, both of which are required for canonical autophagy initiation and autophagosome formation. Furthermore, Tantalosin-induced LC3 lipidation is not sensitive to autophagosome-lysosome inhibition, suggesting that LC3 is not conjugated to autophagosomes and serves a lysosome-independent function. Non-canonical LC3 lipidation has been shown to be involved in LC3-associated phagocytosis (LAP) and LC3-assosiated endocytosis (LANDO) ^23,26,60^. Rubicon is a negative regulator of autophagy and was found to be essential for both LAP and LANDO^50^. However, Rubicon is dispensable for Tantalosin-induced LC3 lipidation, suggesting that Tantalosin induces a process distinct from LAP and LANDO. V-ATPase serves as a common regulator for non-canonical LC3 lipidation by recruiting the E3-like ATG5-ATG12-ATG16L1 complex to membranes ^29,30^. Tantalosin-induced LC3 lipidation is also dependent on the V-ATGPase-ATG16L1 axis. Interestingly, IST1 knockdown, known to induce endosomal recycling impairment, fails to elicit LC3 lipidation. This finding suggests that endosomal recycling defect *pe ser* does not induce LC3 lipidation. Tantalosin-induced LC3 lipidation appears to be correlated with aberrant accumulation of IST1 on membranes. The exact function and mechanism of LC3 conjugation to the stalled recycling endosomes remains to be further elucidated.

In summary, we report the first chemical inhibitor targeting the IST1-CHMP1B interaction within the ESCRT-III complex. This pseudo-natural product compound specifically inhibits the IST1-CHMP1 complex-mediated normal-topology membrane scission and induces non-canonical LC3 lipidation on stalled recycling endosomes. Tantalosin will serve as an invaluable tool to dissect ESCRT-III function in distinct membrane processes and facilitate new insights into the function of non-canonical LC3 lipidation.

## Materials and Methods

### Proteome Integral Solubility Alteration (PISA) assay

HEK293T cells were grown in fifteen 25 cm^2^ flasks at 37°C in 5% CO_2_ using Dulbecco’s Modified Eagle Medium (Lonza, USA) supplemented with 10% FBS (Gibco, Thermo Fisher Scientific), 2 mM l-glutamine (Lonza), and 100 units/mL penicillin/streptomycin (Gibco, Thermo Fisher Scientific). After reaching a 70% confluence, flasks were treated for one hour with 4 µM of Tantalosin, 4 µM of the inactive compound, or the same volume of DMSO. For each condition, five flasks were treated, each representing a biological / experimental replicate of each cell culture treatment. A pool composed by the fifteenth part of each flask was made and processed according to protocols including detergents for stronger protein extraction, and not to the PISA assay, to provide a 16^th^ sample included as carrier proteome in the TMTpro 16-plex for improving detection and quantification of low abundant peptides and proteins across the fifteen PISA samples.

All PISA samples were processed together, according to the published protocol ^38^optimized for higher proteome depth and 16-plex multiplicity using Tandem Mass Tag (TMT) pro (ThermoFisher) and using a gradient for thermal denaturation as described previously^61,62^.

The pool sample was lysed using RIPA buffer complemented with protease inhibitors (Halt protease inhibitors, ThermoFisher), pipetted 10 times, vortexed for 10 s, snap frozen in Liquid N_2_ and then thawed at 37°C. The last two steps were repeated twice, then samples were probe sonicated for 10 cycles of 3 s at 30% amplitude with 3 s of pause between them.

Sample processing for proteomics, based on nanoscale liquid chromatography and tandem mass spectrometry (nLC-MS/MS), was performed according to the PISA assay. The protein concentration of the pool sample and of each soluble fraction after ultracentrifugation of PISA samples was measured using micro-BCA kit (Thermo Fisher) and 50 μg of each sample further processed.

Samples were reduced, alkylated and precipitated using cold acetone, according to previous protocols^38,61^. Proteins pellets were then digested using first LysC and then trypsin enzymes, then digested samples were labelled using the TMTpro 16-plex reagent kit (Thermo Fisher). The final 16-plex sample was cleaned and desalted, and peptides were separated by reversed phase chromatography at high pH, then collected into 48 fractions using a capillary UPLC system. nLC-MS/MS analysis was performed on all 48 fractions using 95-minute gradient per run (total run time 120 minutes per sample) with a system composed by a nano-LC Ultimate 3000 connected to a Orbitrap Q-Exactive HF mass spectrometer (Thermo Scientific).

The database search of nLC-MS fractions for protein identification and quantification was carried out using MaxQuant software v1.6.2.3 (Max Planck Institute of Biochemistry)^63^ against the complete UniProt human proteome database (UP000005640). A 1% false discovery rate was used as a filter at both protein and peptide levels. After removing contaminants, only proteins with at least two unique peptides were included in the final dataset. The TMT reporter ion intensity of each protein in each PISA sample of the 16-plex was normalized by the total ion abundance of the TMT reporter relative to that sample, and then normalized by the average value of the DMSO controls. The statistical significance of variation (P-value) of the soluble amount variation of each protein for either Tantalosin or the inactive compound was calculated by student’s t-test across all five biological replicates in respect to DMSO controls. The PISA results of the Tantalosin’s treatment were also calculated in respect to the inactive compound.

### HDX-MS

3 μL of 70 μM IST1 was mixed with 7 μL of 10 μM Tantalosin (21 μM of IST1 and 7 μM of Tantalosin in the mixture) and incubated at 4°C overnight. 1% DMSO was used as control instead of Tantalosin. 4 μL incubation solution was mixed with 24 μL of stock buffer (50 mM Tris at pH 8.0 including 300 mM NaCl) in D_2_O for 15 s, 150 s, 600 s and 1500 s. After adding 25 μL of ice-cold 100 mM phosphate pH 2.3 and 3.3 M urea, the samples were analyzed in a semi-automated HDX-MS system in which manually injected samples were automatically digested, cleaned and separated at 4°C. Samples were digested using enzymeate BEH pepsin column (WATERS) followed by a 6-min desalting step using 0.2% FA at 60 μL/min. Peptic peptides were then separated by a 21 mm I. D × 50-mm length C18/3 μm column (Thermo Fisher Scientific) using a 20 min/5%–90% acetonitrile gradient in 0.2% formic acid at 60 μL/min. An Orbitrap Q-Exactive mass spectrometer (Thermo Fisher Scientific) operated at 60,000 resolution at m/z 200 was used for analysis. The HDExaminer software (Sierra Analytics, USA) was used to process all HDX-MS data.

### Plasmids, protein expression and purification

The plasmid mCherry-SopF was a gift from Prof. Feng Shao^64^. CHMP2AΔC9-161, CHMP3 and VPS4A were gifts from Lars-Anders Carlson (Umeå University, Umeå, Sweden). The plasmids of pET16b IST1 1-366 and pGEX2T CHMP1B 1-196 were purchased from DNASU (HsCD00671538 and HsCD00520966). His-MBP-TEV tagged IST1 (or CHMP1B) was generated by subcloning IST1 (or CHMP1B) into the pMAL-His-MBP-TEV vector using NdeI/BamHI restriction sites for protein expression and purification. IST1 mutants (IST1^I141D^ and IST1^G142D^) were generated from wild-type IST1 via PCR mutagenesis.

Full length of IST1 and CHMP1B were expressed in *E.coli* strain BL21(DE3) cells and grown in LB broth supplemented with 100 mg/L ampicillin (4 liters of cultures for each protein). Cells were grown at 37°C on a shaker (160 rpm) until the absorbance at 600 nm (OD600) reached 0.5-0.7. The protein expression was induced with 0.2 mM Isopropyl β-d-1-thiogalactopyranoside (IPTG) and incubate cells at 20°C (150 rpm) overnight (< 18 h). The cells were harvested by centrifugation at 5000 rpm, 4°C for 20 min, then discard the supernatant carefully and wash the cells with PBS by centrifugation.

All purification steps were performed at 4°C unless otherwise noted. Cells were resuspended in lysis buffer (50 mM Tris, 300 mM NaCl, 5% glycerol, pH 7.5) freshly added 1 mM phenylmethylsulfonyl fluoride (PMSF) and 2 mM β-mercaptoethanol. Resuspended cells were lysed by cell disruptor (Constant Systems) at 27 kpsi for three times. The cell lysates were further supplemented with 1% Triton X-100 and then centrifuged at 22000 rpm for 1 h. The supernatant was filtered through a 0.2 µm membrane and then loaded onto a 5 ml HisTrap HP (Cytiva) column, which was equilibrated with buffer A (50 mM Tris, 300 mM NaCl, 5% glycerol and freshly added 2 mM β-mercaptoethanol, pH 7.5) and then protein was eluted with an isocratic of buffer B (50 mM Tris, 300 mM NaCl, 5% glycerol, 500 mM Imidazole and fresh 2 mM β-mercaptoethanol, pH 7.5). The eluted protein was incubated with 1%(w/w) TEV protease and dialyzed against cleavage buffer (50 mM Tris, 250 mM NaCl, 5% glycerol, fresh 2 mM β-mercaptoethanol, pH 8.0) overnight to remove the His-tag. The cleaved protein was purified on HisTrap HP (Cytiva) column again and the flow through fractions were collected. The fraction of interest was concentrated with 10kDa Amicon Ultra centrifuge filter to ∼10 ml. Finally, the protein was purified by size-exclusion chromatography column (Superdex 200 pg, HiLoad 26/60) in 50 mM Tris, 300 mM NaCl, 5% Glycerol and fresh 2 mM β-mercaptoethanol, pH 8.0. Fractions containing the peak of interest were concentrated with Amicon 10 kDa to a final concentration of ∼8.9 mg/mL (IST1 wild-type) ∼17.22 mg/mL (IST1_I141D_), ∼13.52 mg/mL (IST1_G142D_) and ∼9.2 mg/ml (CHMP1B). 2 mM TCEP were added to proteins for storage.

### Synthesis of MIM1 peptide

CHMP1B C-terminal peptide MIM1(CHMP1B_183-199_) was synthesized based on Fmoc-based solid phase peptide (SPPS) synthesis strategy. Unless otherwise noted, all of the natural amino acids used for coupling were standard Fmoc protected L-amino acids. All peptides were synthesized using peptide synthesizer Syro I (Biotage, Sweden). Briefly, the resin (Rink Amide ChemMatrix resin, Biotage) was swelled in DMF for 1 h. Next, the subsequent N-terminal peptide chain elongation was achieved by standard Fmos strategy employing the commercially available amino acid building blocks Fmoc-Ala-OH, Fmoc-Asp(OtBu)-OH, Fmoc-Glu(OtBu)-OH, Fmoc-Leu-OH, Fmoc-Gln(Trt)-OH, Fmoc-Arg(Pbf)-OH, Fmoc-Ser(tBu)-OH and Fmoc-Val-OH. Fmoc group was cleaved by shaking the resin twice for 10 min with a solution of piperidine/DMF (1:4, v/v). The coupling was performed for 2 h (4.0 eq. amino acid acid, 4.0 eq. HBTU and 2.0 eq. HOBT, 4.0 eq DIPEA) in DMF. After synthesis, the resin was washed thoroughly with DMF and DCM, then dried under vacuum overnight. Next, the resin was treated with a solution TFA/H_2_O/TIPS (95:2.5:2.5, v/v/v) at r.t. for 2 h to release the peptide. Then the resin was filtered and rinsed twice with TFA. The crude peptide was obtained by precipitation with adding cold diethyl ether. The crude peptide was purified by preparative HPLC. And masses were confirmed either before labeling or after labeling by mass spectrometry.

### Fluorescent labeling of IST1

For the FITC labeling, IST1 protein (100 μM) was incubated with 3 equivalents FITC (300 μM) in 50 mM sodium phosphate buffer (pH 8.0, including 300 mM NaCl) for 3 h at room temperature. Labeled protein was separated from free fluorescent dye using a gel filtration column (GE Healthcare illustra NAP-10 Columns Sephadex G-25), followed by ESI-MS analysis (Agilent 6230 TOF LC/MS) and SDS-PAGE analysis. The degree of labeling (DOL) is calculated from following equations, CF = A280 free dye/Amax free dye, DOL = Amax × ε280/ ([A280–Amax × CF] × ε578). A280 is the absorbance of the protein–dye conjugate at 280 nm; A578 Amax is the absorbance of the protein–dye conjugate at its maximum absorbance; ε280 is the extinction coefficient of the protein at 280 nm in cm^−1^ M^−1^; εdye is the extinction coefficient of the dye at its maximum absorbance in cm^−1^ M^−1^; and CF is the correction factor and CF values for FITC is 0.3. The DOL of FITC-labeled IST1 is 0.41.

### Fluorescent labeling of MIM1

FAM labeled MIM1 was synthesized by using solid phase peptide synthesis method and purified on HPLC. Briefly, 25 mg of MIM1 resin was resuspended in 2 mL DMF solvent and swelled for 1 h. Next, add 19.7 mg FAM (5.0 eq), HOBt (4.0 eq, 7.0mg), EDC (10 mg) in 1 mL DMF solvent to the resin, leave the reaction on a shaker with continuously shaking at room temperature for overnight. After synthesis, the resin was washed thoroughly with DMF and DCM for twice, then dried under vacuum overnight. Next, the resin was treated with a solution TFA/H_2_O/TIPS (95:2.5:2.5, v/v/v) at rt for 2 h to release the labelled peptide. Then the resin was filtered and rinsed twice with TFA. The crude FAM labeled peptide was obtained by precipitation with adding cold diethyl ether. The crude peptide was purified by preparative HPLC. And masses were confirmed either before labeling or after labeling by mass spectrometry.

### MST measurement

Binding affinity measurement of proteins was performed using a Nanotemper Monolith NT.115 instrument. MST experiments were performed in assay buffer (50 mM Tris, pH 8.0, 150 mM NaCl, 0.05% Tween 20 and 1 mM dithiothreitol (DTT)) using 80 nM FITC-labeled IST1 or 50 nM FAM-labeled MIM1 mixed with different concentrations of proteins in MST assay buffer. The capillary was excited at 480 nm. Dissociation constants were calculated by fitting the increase in to a 1:1 binding equation using Nanotemper Monolith MO. Affinity software. For Tantalosin treatment, IST1 was pre-treated with tantalosin (final concentration is 7 µM) at room temperature for 4 h. Experiments were performed at 22°C, and samples were filled into standard capillaries for measurement. Each binding was measured at least three times independently, and mean K_D_ values were reported ± SD.

### TEM sample preparation and imaging

Membrane-free IST1 and CHMP1B polymers were prepared according to the protocol described by McCullough et al^18^, but with modified protocol to meet the experiment requirement. Briefly, IST1and CHMP1B (32 μM final protein concentrations) in 50 mM Tris, pH 8.0, 300 mM NaCl, 5% (w/v) glycerol, 5 mM BME were serial diluted in dilution buffer (50 mM Tris, pH 8.0) with a final salt concentration of 25 mM NaCl. Reactions were then incubated at room temperature overnight. The assemblies were concentrated by low-speed centrifugation (12700 rpm, 20 min) and re-suspended in 10 μl low ionic strength buffer (50 mM Tris, pH 8.0, 25 mM NaCl).

CHMP2A-CHMP3 tubes were prepared according to the method of Lata et. al^40^. 15 μM MBP tagged CHMP2AΔC and 30 μM CHMP3 were mixed and incubated together in HBS (20 mM HEPES, pH 7.6, 150 mM NaCl) buffer for overnight. The assemblies were concentrated at 12700 rpm for 20 min and re-suspended in 100 μL HBS buffer.

For negative staining, 3 µl of the co-polymer was applied to a continuous carbon grid (Ted Pella, 300 mesh), it was stained three times for 10 s with drops of 1.5% uranyl acetate, blotted, and allowed to air dry. Imaging was performed at a Thermo Fisher Talos L120C operated at 120 kV, equipped with a 4096-× 4096-pixel Ceta camera. The nominal magnification was 28000× pixel size. The length of the filaments was measured using ImageJ and plotted using GraphPad Prism v10.0.1.

### Sedimentation assay

IST1 and CHMP1B, or MBP-CHMP2AΔC and CHMP3 proteins were incubated with vehicle (DMSO), inactive compound (7 μM) or Tantalosin (7 μM) at room temperature for 12 h. The mixtures of protein were centrifuged at 4°C, 12,700 rpm for 1 hour. Equal amounts of samples from pellet and supernatant were heated at 95°C with 1X Laemmli buffer and separated by SDS-PAGE. Gels were stained with Coomassie blue and imaged with a Bio-rad Chemidoc. Images were analyzed by ImageJ software.

### Antibodies and reagents

The following antibodies were used in this study. Immunofluorescence: anti-IST1 (1:100, 51002-1-AP, Proteintech), anti-Transferrin Receptor (1:100, 13-6800, Invitrogen), anti-LC3B (1:500, PM036, MBL International), anti-LC3B (1:50, M152-3, MBL International), anti-α-Tubulin (1:1000, T6199, Sigma-Aldrich), anti-EGFR (1:100, #4267, Cell Signaling Technology), anti-GFP (1:500, A10262, Invitrogen). Secondary antibodies were goat anti-rabbit/mouse Alexa Fluor 488 conjugated, goat anti-rabbit/mouse Alexa Fluor 568 conjugated, goat anti-rabbit Alexa Fluor 647 conjugated, goat anti-mouse DyLight 649 conjugated, and all were purchase from Thermo Fisher Scientific.

Western Blot: anti-IST1 (1:1000, PA5-62981, Invitrogen), anti-Transferrin Receptor (1:1000, 13-6800, Invitrogen), anti-LC3B (1:1000, #2775, Cell Signaling Technology), anti-ATG7 (1:1000, #8558, Cell Signaling Technology), anti-ATG16L1 (1:1000, #8089, Cell Signaling Technology), anti-FIP200 (1:1000, #12436, Cell Signaling Technology), anti-ATG9A (1:1000, #13509, Cell Signaling Technology), anti-Rubicon (1:1000, #8465, Cell Signaling Technology), anti-EGFR (1:100, #4267, Cell

Signaling Technology), anti-ALIX (1:1000, #634502, BioLegend), anti-TSG101 (1:1000, #HPA006161, Sigma-Aldrich), anti-actin (1:10000, A2228, Sigma-Aldrich), anti-vimentin (1:1000, #5741, Cell Signaling Technology), anti-β-tubulin (1:1000, MA5-16308, Invitrogen). Secondary antibodies were goat anti-rabbit/ mouse HRP conjugated (31460/ 31430, 1:10000, Thermo Fisher Scientific).

Chloroquine (C6628) was purchased from Sigma Aldrich, Transferrin from Human Serum, Alexa Fluor™ 555 Conjugate (T35352) was purchased from Thermo Fisher Scientific.

### Cell culture

HeLa (ATCC), HEK293T (ATCC), MCF7 cells stably expressing EGFP^35^, HeLa (WT, ATG7 KO, FIP200 KO, ATG9A KO, Rubicon KO, ATG16 L1 KO - kind gift of Tamotsu Yoshimori, Osaka University, Osaka, Japan ^65^) were cultured in Dulbecco’s Modified Eagle’s Medium, (DMEM, Sigma Aldrich) supplemented with 10% fetal bovine serum (FBS, Thermo Fisher Scientific,), non-essential amino acids (NEAA, Thermo Fisher Scientific) and penicillin/streptomycin (Pen/Strep, Thermo Fisher Scientific). For imaging purposes cell were cultured in phenol-free DMEM (Sigma Aldrich) supplemented with 10% FBS, NEAA, Pen/Strep and 20 mM HEPES (Thermo Fisher Scientific). Cells were routinely tested for the mycoplasma absence using the LookOut Mycoplasma PCR detection kit (Sigma-Aldrich).

### Plasmids and transfection

Transfection was performed using X-tremeGENE™ HP DNA Transfection Reagent (Roche) in Opti-MEM™ medium (Thermo Fisher Scientific) according to manufacturer’s protocol. siRNA for IST1 was designed and purchased from ThermoFisher Scientific (1 – ID: s18939, 2 – ID: s18940, 3 – ID: S18941). siRNA transfection was performed using RNAiMAX reagent (13778150, Thermo Fisher) according to manufacturer’s instructions.

### Transferrin pulse-chase assay

Cells were treated with serum-free media at 37 °C for 1 hour in the presence or absence of Tantalosin to hold receptors at the plasma membrane. Then 25 μg/ mL of Transferrin from Human Serum, Alexa Fluor™ 555 Conjugate (T35352, Invitrogen) was added to media, and cells were placed on ice for 30 min. After quick wash, media was changed to the imaging phenol-free media with or without Tantalosin, and live-cell confocal imaging was immediately started. For VEH and Tantalosin-treated cells all imaging conditions were kept identical.

### Evaluation of cytokinesis

For evaluation of cytokinesis HeLa cells were transfected siRNA for 48 h or treated with the compound for 24 h. Immunofluorescence staining was performed using anti-α-Tubulin antibodies and nuclei were stained with DAPI. 120-180 cells were analyzed per replicate and scored for the presence of remaining midbodies.

### EGFR degradation assay

HeLa cells were incubated in DMEM media containing 0.25% FBS for 16 h. Then cells were pre-treated with VEH or 4 μM Tantalosin for 2 h, followed by treatment with 50 ng/ml EGF for 30, 60 or 120 minutes.

### Exosome purification and analysis

HeLa cells were incubated in 10×150 cm flasks per condition in serum-free OPTI-MEM medium supplemented with Pen/Strep for 24 h in presence of absence of Tantalosin. All subesquent procedures with the conditioned media from these cells were performed at 4°C. Conditioned media was sequentially centrifugated at 600×g for 10 min, at 2000×g for 10 min and at 10000×g for 30 min. Then pre-cleared media was concentrated using Vivaflow® 50R cassette with 100 kDa molecular cut-off (VF05H4, Sartorius). Concentrated media was centrifugated at 100000×g for 4 h. Pellets of EVs were resuspended in PBS overnight at 4°C, then used for TEM or western blot analysis.

EVs (5 µl, 1-2 µg protein) were added to carbon-coated and glow-discharged copper grids. Grids were incubated for 5 min at RT, Excess solution was blotted and the grids were washed 2 times with milliQ water. Thereafter, the grids were stained with 1.5% uranyl acetate and dried at RT before TEM images were acquired (FEI Talos L 120C).

### Western blot

Cells were washed with PBS and lysed in ice-cold lysis buffer containing 0.25% Nonidet P-40, 20 mM Tris-HCl (pH 8), 300 mM KCl, 10% Glycerol, 0.5 mM EDTA, 0.5 mM EGTA supplemented with protease inhibitor cocktail (Roche) and 1 mM phenylmethylsulfonyl (PMSF). Lysates were incubated 20 min on ice, passed six times through a 21G needle and cleared at 18000×g. The protein concentrations of lysates were determined using Bradford assay (Bio-Rad). Cell lysates were diluted in 1X SDS-sample buffer (Bio-Rad) supplemented with 355 mM β-mercaptoethanol and incubated at 95°C for 10 min. Proteins were separated using SDS-PAGE and transferred to nitrocellulose membrane using Trans-Blot Turbo transfer system (Bio-Rad). Membranes were blocked in 5% skimmed milk diluted in Tris-buffered saline with 0.1% Tween-20 (TBS-T) for 1 hour at RT. Membranes were washed three times with TBS-T and incubated with primary antibodies overnight at 4°C. After washing three times with TBS-T membranes were incubated with secondary HRP-conjugated antibodies for 1 hour at RT. Chemiluminescent signal was developed using Clarity Western ECL Substrate (Bio-Rad) and imaging was performed using ChemiDoc Imaging System (Bio-Rad). Densitometric intensities of bands were quantified using ImageLab software or ImageJ software.

### Immunofluorescence

For immunofluorescence experiments cells were cultured on No. 1.5 glass coverslips. Cells were quickly washed with PBS and fixed with 4% methanol-free paraformaldehyde (Sigma Aldrich) for 10 min at room temperature. Cells were washed three times with 1.5 mg/ml glycine in PBS and permeabilized with 0.25% Triton X-100 for 5 min at room temperature. Cells were washed three times with PBS and incubated in 5% donkey serum (blocking solution) for 30 min at room temperature. Cells were incubated with primary antibodies diluted in 5% donkey serum for 1-2 h at room temperature. Cells were washed 3 times with PBS and incubated with corresponding secondary fluorescently labelled antibodies for 30 min at room temperature. When indicated, cells were stained with DAPI and mounted on imaging slides with ProLong Diamond Mounting Media (Invitrogen).

### Confocal microscopy

Fluorescent microscopy was performed on a Leica SP8 FALCON inverted confocal microscope (Leica Microsystems) equipped with an HC PL APO 63x/1.40 oil immersion objective. For live-cell imaging cell were maintained in temperature-controlled hood at 37 °C and 5% CO_2_. The microscope was operated using Leica Application Suite X (LASX). Fluorophores were excited using a 405 nm Diode or tuned white light lasers, and scanning was performed in line-by-line sequential mode.

### Imaging analysis

Imaging analysis was performed using ImageJ software (NIH). EGFP-LC3 and intracellular transferrin areas were quantified using the “Analyze particles” plugin and normalized to total cell area. Tf-555-assosiated EGFP-LC3 fluorescence was calculated by measuring mean EGFP fluorescence over time within a mask generated using the Alexa-555 channel. Fluorescence intensity profiles of the line scans were determined using the “Plot Profile” analysis function.

### Statistics

Statistical analysis was performed using GraphPad 9.0.0 software. Statistical significance was determined using unpaired two-tailed Student t-test or one-way ANOVA with Tukey test for multiple comparisons. *P<0.01, ** P<0.001, *** P<0.0001, **** P<0.00001; ns – not significant.

## Acknowledgements

This work was supported by the European Research Council (ChemBioAP), Vetenskapsrådet (Nr. 2018-04585, Nr. 2022-02932), the Knut and Alice Wallenberg Foundation and the Göran Gustafsson Foundation for Research in Natural Sciences and Medicine to Y.W.W. We acknowledge the Biochemical Imaging Center at Umeå University and the National Microscopy Infrastructure, NMI (VR-RFI 2019-00217) for providing assistance in microscopy.

## Author contributions

AK – investigation, formal analysis, data curation, visualization, writing – original draft, writing – review and editing; SL – investigation, formal analysis, data curation, writing – original draft, writing – review and editing, DPC – investigation, formal analysis, writing – review and editing; KS, LKH, XC, BS, GN – investigation, writing – review and editing; MG - investigation, supervision, writing – review and editing; JDG, LAC, SZ – supervision, writing – review and editing, HW – conceptualization, supervision, writing – review and editing; YW - conceptualization, funding acquisition, project administration, supervision, writing – review and editing.

---

**Supplementary Figure 1.**
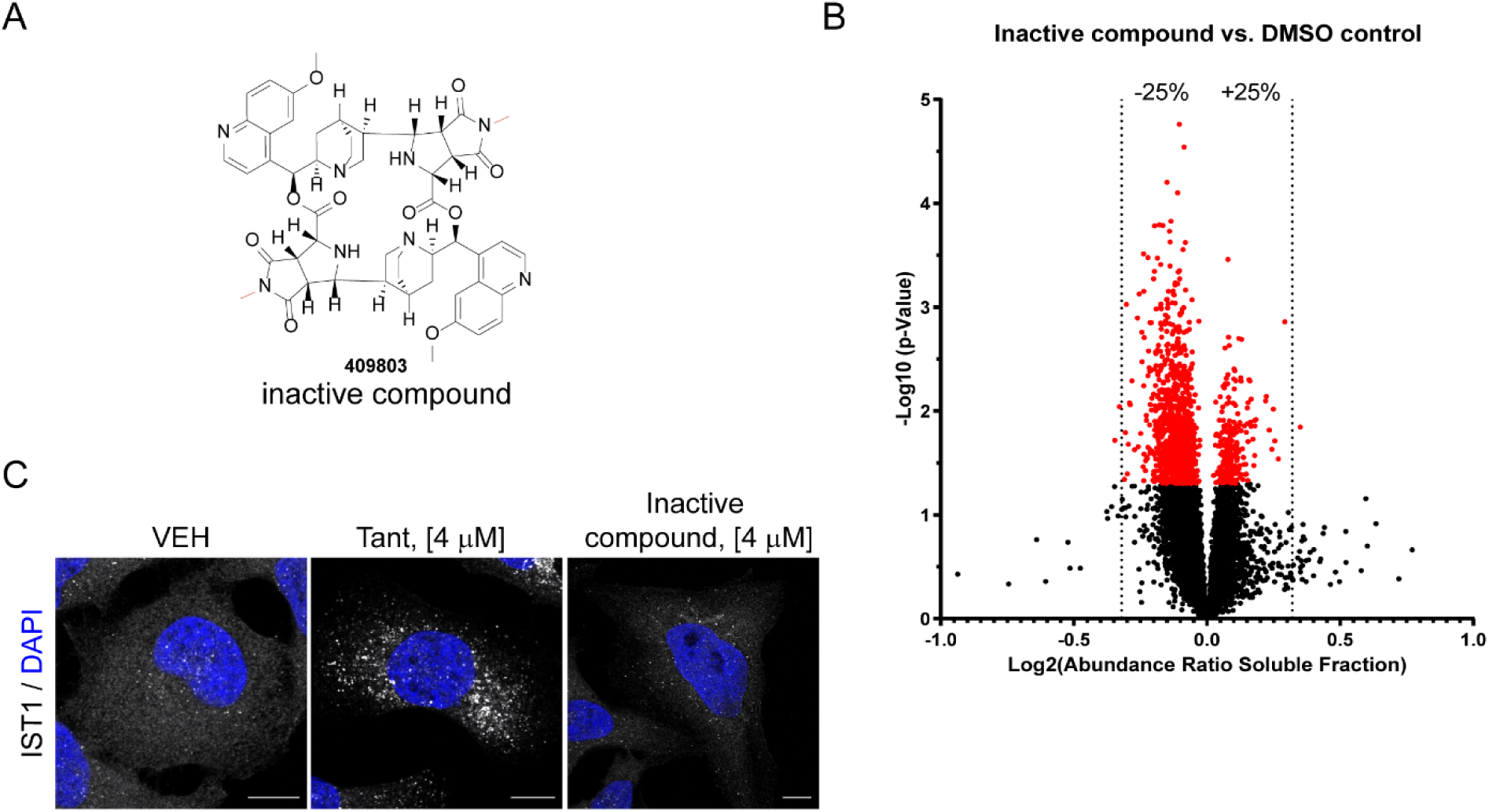
**(A)** Chemical structure of inactive compound. **(B)** Volcano plots of results obtained from the PISA assay performed in HEK 293T cells treated with 4 µM inactive compound versus DMSO-treated cells. **(C)** HeLa cells were treated with vehicle (VEH), 4 µM Tantalosin or 4 µM inactive compound for 4 h and immunostained for endogenous IST1. Nuclei were counterstained with DAPI. Scale bar: 10 µm. Results are representative of three independent experiments.

**Supplementary Figure 2.**
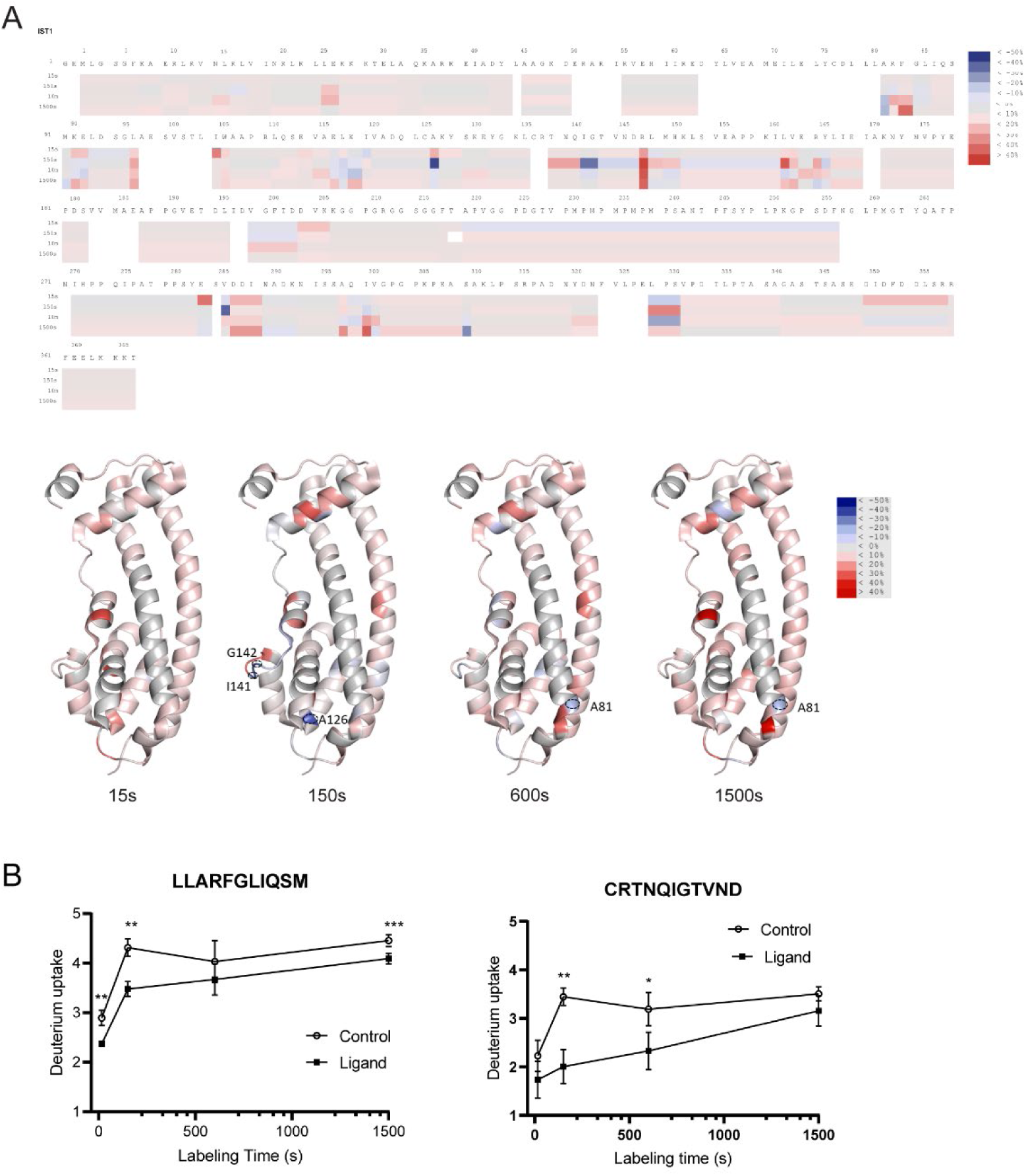
Hydrogen Deuterium Exchange Mass Spectrometry (HDX-MS) analysis of the Tantalosin-IST1 binding. **(A) Upper panel:** Heatmap of HDX-MS analysis identified two possible Tantalosin-binding sites in IST1. Heatmap shows the relative fractional deuterium uptake of IST1 in the presence of Tantalosin as compared to IST1 in the presence of equivalent amounts of DMSO. Experiments were performed at the following exposure time: 15 s, 150 s, 600 s and 1500 s. Coloring is according to the spectrum bar. Residues without coverage are colored in white. **Lower panel:** the relative fractional deuterium assessed by HDX-MS experiments mapped onto the IST1 structure (PDB: 3FRR). Coloring is according to the spectrum bar. **(B)** Comparison of deuterium uptake in peptide 79-89 with and without Tantalosin (left) and comparison of deuterium uptake in peptide 136-146 with and without Tantalosin (right) for three independent experiments described in A. * P<0.01, ** P<0.001.

**Supplementary Figure 3.**
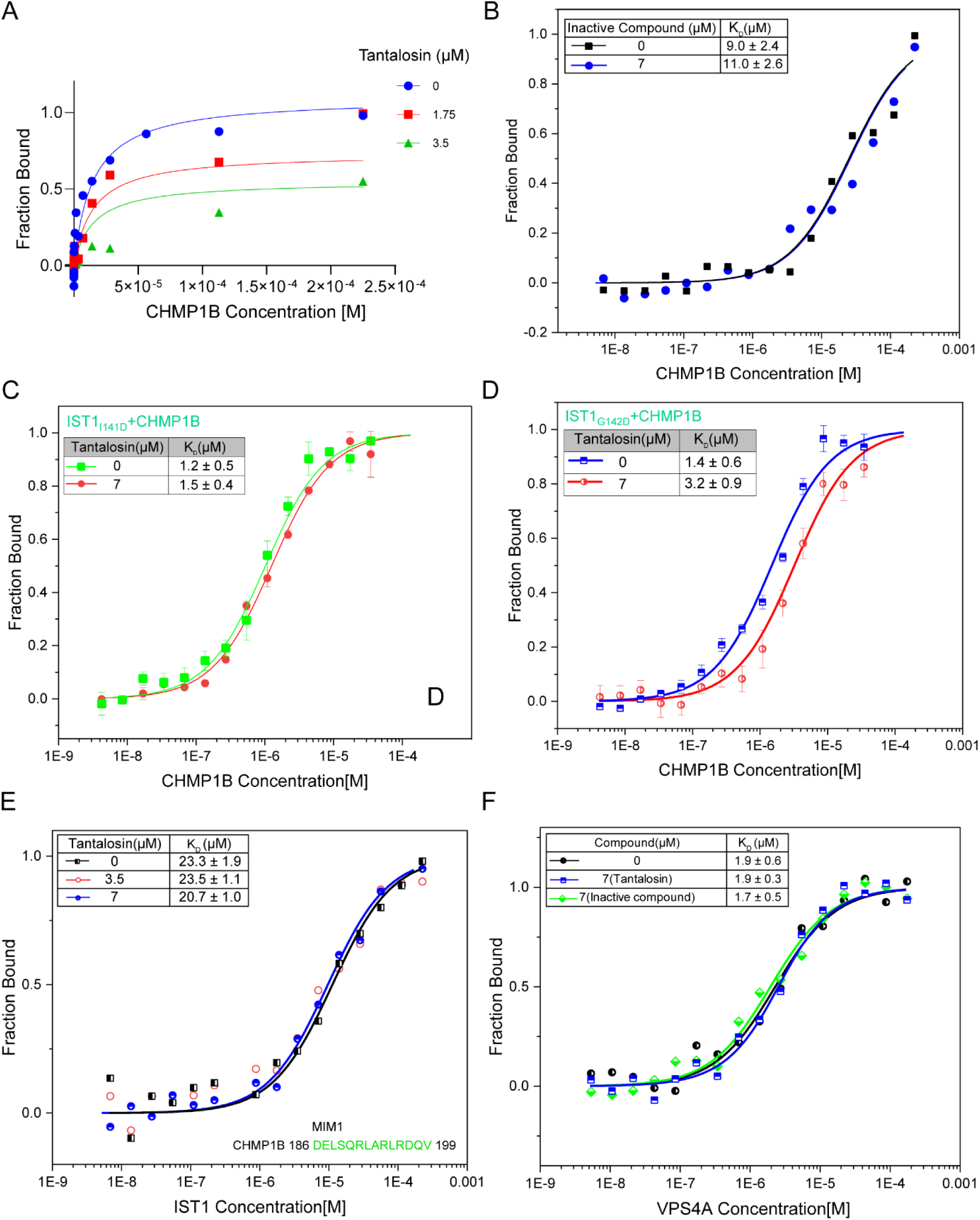
(**A**) Global fitting of the IST1-CHMP1B binding in the presence of indicated concentrations of Tantalosin using a non-competitive inhibition model. **(B)** Dose-dependent binding curves of IST1 and CHMP1B in solution in the presence of indicated concentrations of inactive compound. CHMP1B (225 µM to 6.8 nM) was mixed with IST1 (80 nM) in assay buffer. Inset represents a table of mean K_D_ ± SD for the IST1-CHMP1B interaction from three independent experiments. **(C)** and **(D)** MST analysis of IST1^I141D^ and IST1^G142D^ interaction with CHMP1B in solution. CHMP1B (139 μM to 4.2 nM) was mixed with IST1 mutants (80 nM) in assay. Insets represents mean K_D_ ± SD from three biological replicates. **(E)** MST analysis of IST1 interaction with CHMP1B MIM1 in solution. IST1 (175 μM to 5.3 nM) was mixed with FAM-labeled MIM1 peptide (50 nM) in assay buffer. Insets represents mean K_D_ ± SD (n=3). The sequence of MIM1 is shown in green as an inset. **(F)** MST analysis of IST1 interaction with VPS4A in solution. VPS4A (170 μM to 5 nM) was mixed with IST1 (80 nM) in assay buffer. Insets represents mean K_D_ ± SD (n=3).

**Supplementary figure 4.**
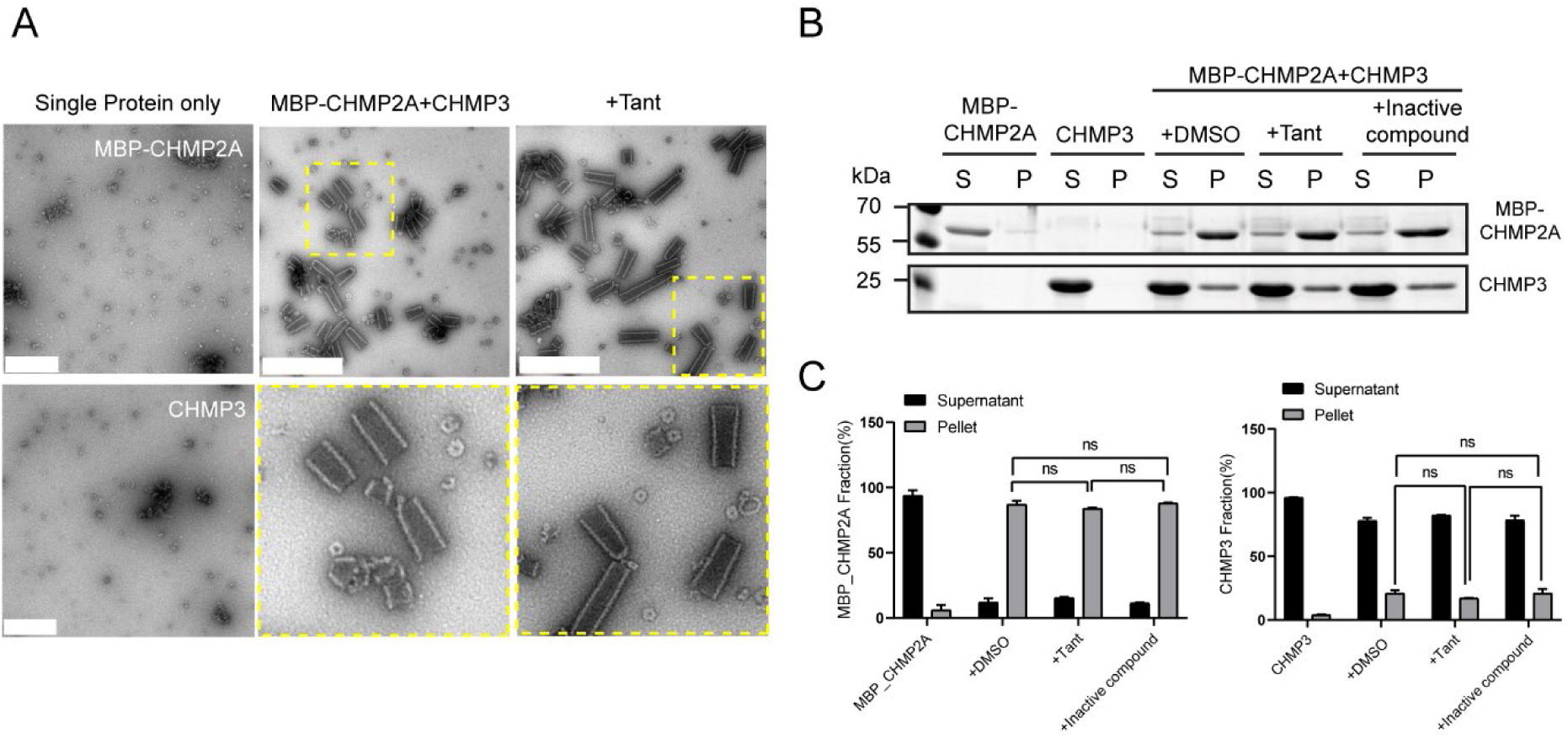
**(A)** TEM images of CHMP2AΔC (with MBP tag) and CHMP3 alone, CHMP3-CHMP2AΔC (with MBP tag) copolymer with or without Tantalosin treatment (7 µM). Scale bar: 0.5 µm. **(B)** SDS-PAGE analysis of MBP-CHMP2AΔC (15 µM) and CHMP3 (30 µM) co-polymer sedimentation with DMSO, Tantalosin (7µM) or inactive compound (7 µM). S: supernatant; P: pellet. **(C)** Quantification of CHMP2A and CHMP3 levels for experiment described in (B). Statistical analysis was determined from three biological replicates using a two-tailed, unpaired t-test. ns: not significant.

**Supplementary Figure 5.**
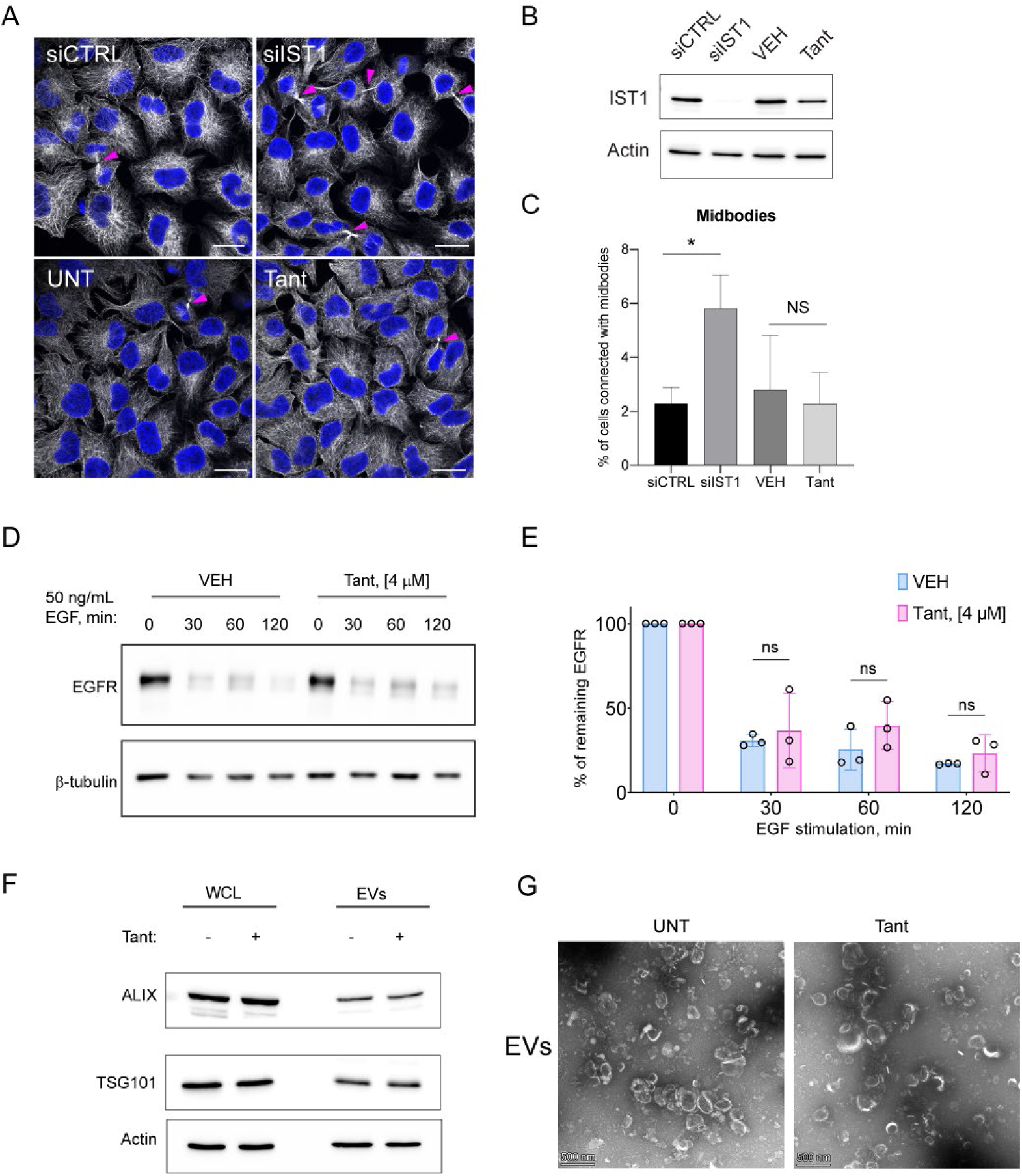
Tantalosin does not affect cytokinesis, MVB sorting and exosome biogenesis. **(A)** Representative images of HeLa cells transfected with control siRNA (siCTRL) or siIST1 for 48 h, and treated with VEH or 1 μM Tantalosin for 24 h. Cells were immunostained for α-Tubulin, nuclei were counterstained with DAPI. Magenta arrowheads indicate midbodies. Scale bar: 20 µm. **(B)** Western blot analysis of IST1 levels in cells treated as indicated in (A). Actin serves as a loading control. **(C)** Quantification of the percentage of cells with arrested midbodies (left panel) following siRNA or Tantalosin treatment as mentioned in (A). n = 3 independent experiments, *P < 0.05, NS – not significant. **(D)** Western Blot analysis of EGFR levels in HeLa cells stimulated with 50 ng/ml EGF for 30, 60, 120 minutes. Prior to EGF cells were treated with VEH or 4 μM Tantalosin for 2 h. **(E)** Quantification of EGFR levels upon EGF stimulation corresponding to (D). Percentage of the remaining EGFR is calculated relative to 0 min EGF stimulation for each of VEH or Tantalosin-treated cells. Bar graphs represent mean ± SD for n = 3 independent experiments, which are indicated as points. Statistical analysis is performed for each time point of EGF stimulation for the comparison of VEH and Tantalosin treatment using two-tailed Student t-test; ns: not significant. **(F)** Western Blot analysis of TSG101 and ALIX levels in whole cell lysate (WCL) or extracellular vesicles (EVs) purified from conditioned media collected from cells treated with VEH or 1 μM Tantalosin for 24 h. **(G)** Representative transmission electron microscopy (TEM) images of EVs purified from conditioned media collected from cells treated with VEH or 1 μM Tantalosin for 24 h. Scale bar: 500 nm.

**Supplementary Figure 6.**
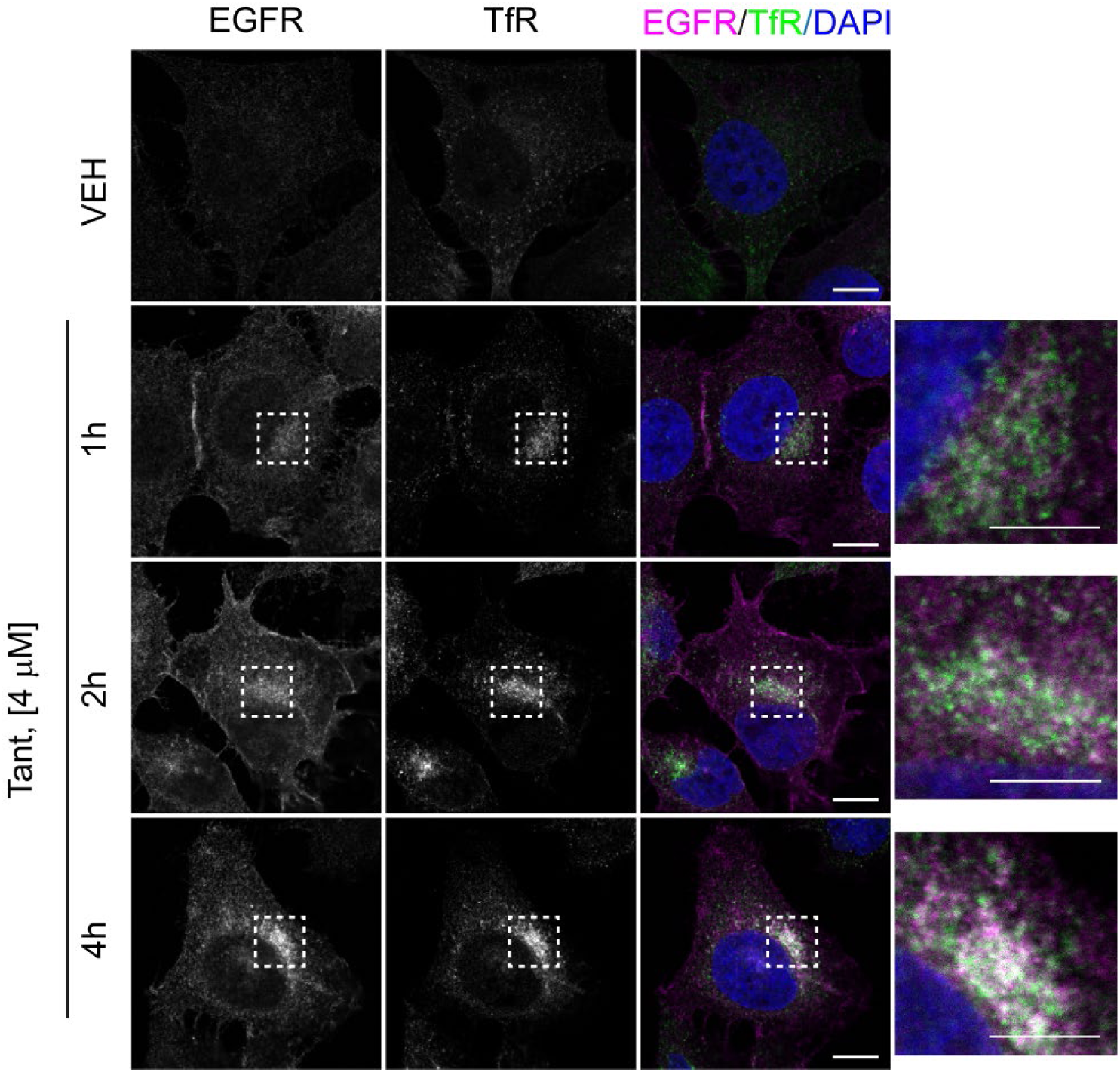
Representative images of HeLa cells treated with 4 µM Tantalosin for 1, 2 and 4 h and immunostained for EGFR and transferrin receptor (TfR). Scale bar: 10 μm for original images and 5 μm for insets.

**Supplementary figure 7.**
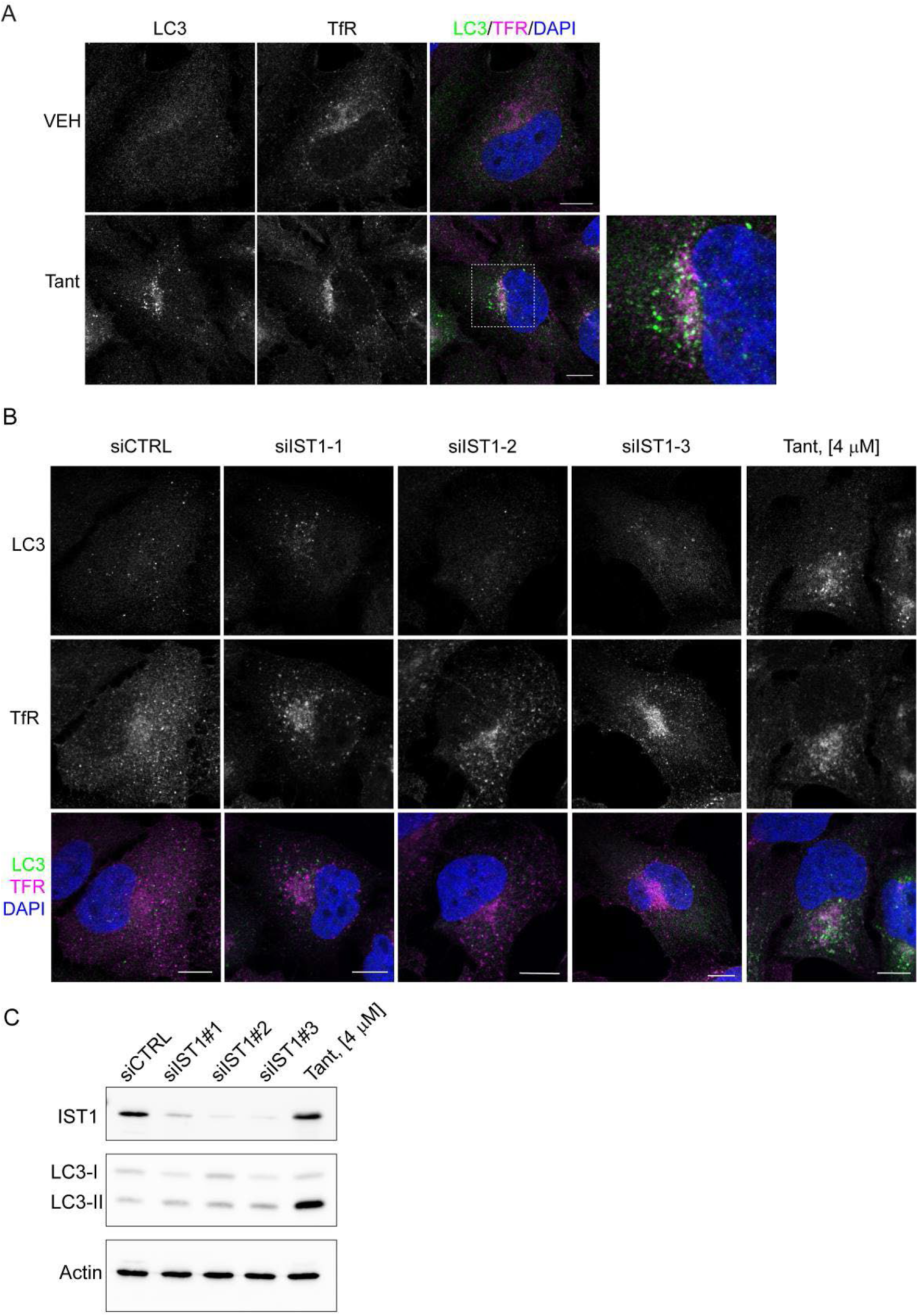
**(A)** Representative images of HeLa cells treated with VEH or 4 µM Tantalosin for 2 h and immunostained for endogenous LC3 and TfR. Scale bar: 10 μm. **(B)** Representative confocal images of HeLa cells transfected with control siRNA, three individual siRNAs for IST1 knockdown or treated with 4 µM Tantalosin for 4 h. Cells were immunostained for LC3 and transferrin receptor (TfR). Scale bars: 10 μm. **(C)** Western blot analysis of IST1 and LC3 levels in cell transfected with control siRNA, three individual siRNAs for IST1 knockdown or treated with 4 µM Tantalosin for 4 h. Actin serves as a loading control.

**Supplementary Figure 8.**
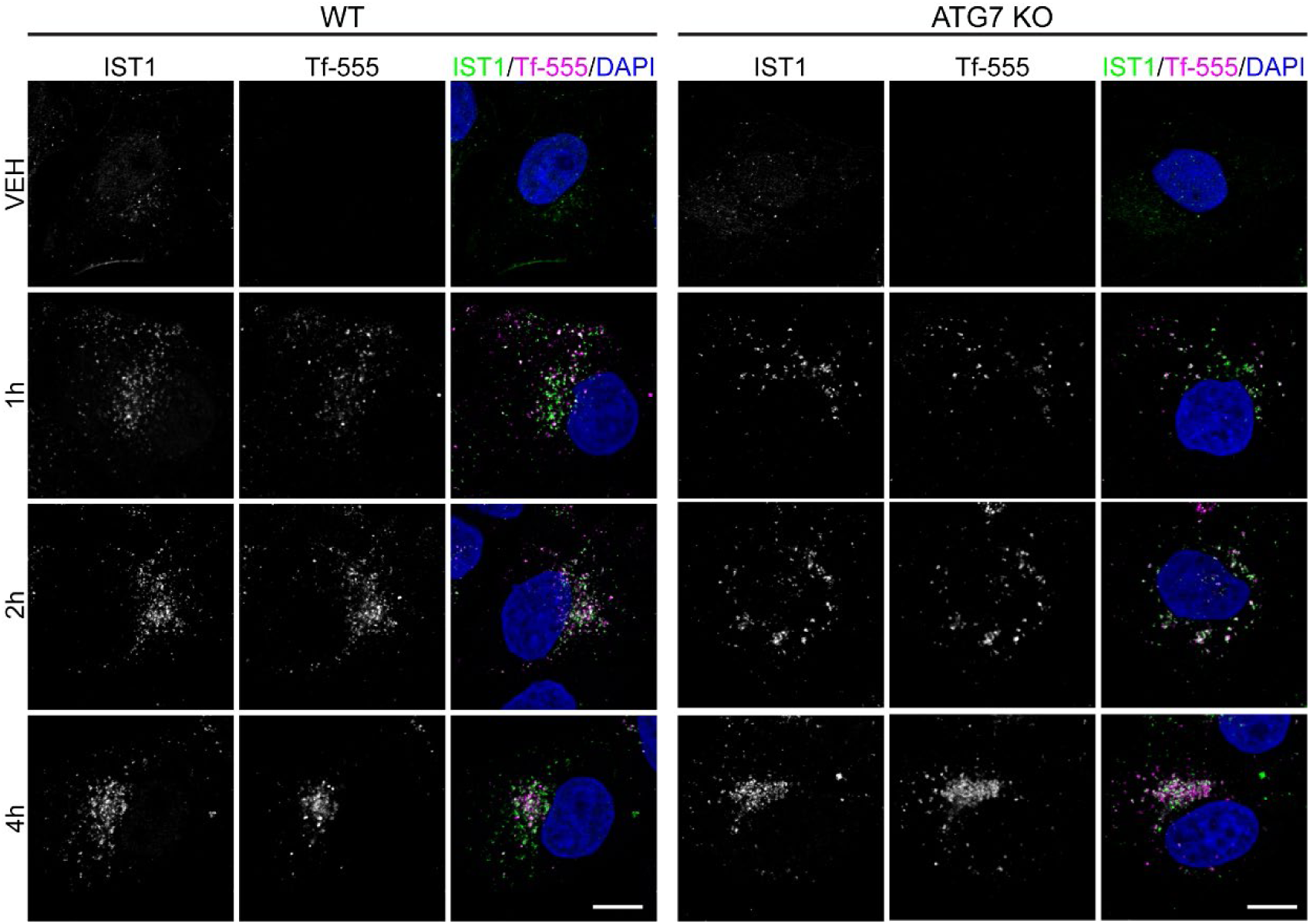
Representative images of WT or ATG7 knock-out (ATG7 KO) HeLa cells treated with fluorescently labelled transferrin (Tf-555) and 4 µM Tantalosin for indicated time and immunostained for IST1. All images were acquired with the same size, scale bars: 10 μm.

**Supplementary Figure 9.**
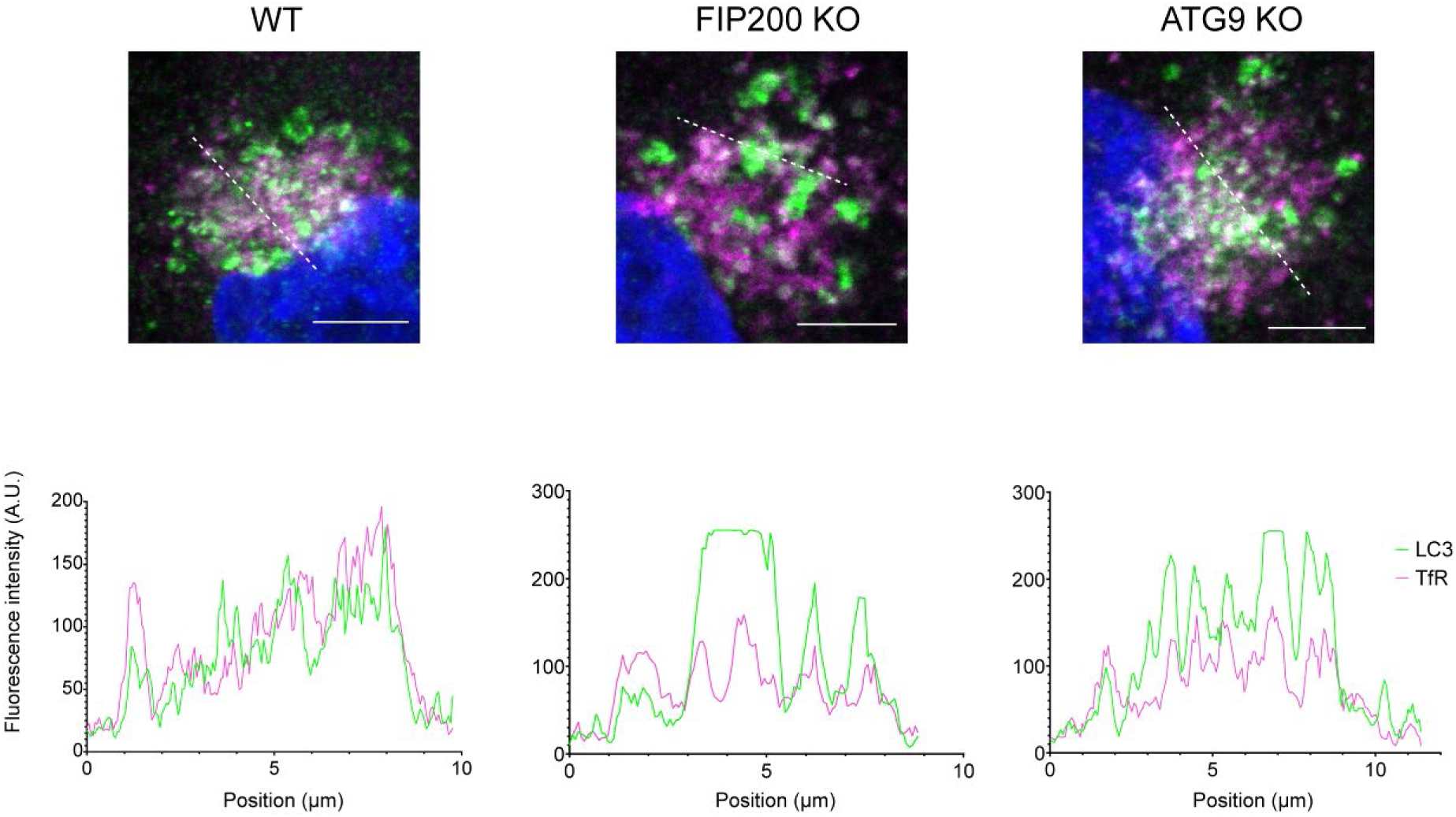
Fluorescent intensity profiles for LC3 and TfR immunostaining for WT, FIP200 KO and ATG9 KO cells presented in Figure 5.

## References

1. Vietri, M., Radulovic, M. & Stenmark, H. The many functions of ESCRTs. Nat Rev Mol Cell Biol 21, 25–42 (2020).

2. Katzmann, D. J., Babst, M. & Emr, S. D. Ubiquitin-dependent sorting into the multivesicular body pathway requires the function of a conserved endosomal protein sorting complex, ESCRT-I. Cell 106, 145–55 (2001).

3. Babst, M., Katzmann, D. J., Snyder, W. B., Wendland, B. & Emr, S. D. Endosome-associated complex, ESCRT-II, recruits transport machinery for protein sorting at the multivesicular body. Dev Cell 3, 283–289 (2002).

4. Colombo, M. et al. Analysis of ESCRT functions in exosome biogenesis, composition and secretion highlights the heterogeneity of extracellular vesicles. J Cell Sci (2013) doi:10.1242/jcs.128868.

5. Takahashi, Y. et al. An autophagy assay reveals the ESCRT-III component CHMP2A as a regulator of phagophore closure. Nat Commun 9, 1–13 (2018).

6. Zhou, F. et al. Rab5-dependent autophagosome closure by ESCRT. J Cell Biol 218, 1908–1927 (2019).

7. Feng, Q. et al. MAPT/Tau accumulation represses autophagy flux by disrupting IST1-regulated ESCRT-III complex formation: a vicious cycle in Alzheimer neurodegeneration. Autophagy 16, 641–658 (2020).

8. Votteler, J. & Sundquist, W. I. Virus budding and the ESCRT pathway. Cell Host Microbe 14, 232–41 (2013).

9. Carlton, J. G. & Martin-Serrano, J. Parallels between cytokinesis and retroviral budding: a role for the ESCRT machinery. Science 316, 1908–12 (2007).

10. Vietri, M. et al. Spastin and ESCRT-III coordinate mitotic spindle disassembly and nuclear envelope sealing. Nature 522, 231–5 (2015).

11. Olmos, Y., Hodgson, L., Mantell, J., Verkade, P. & Carlton, J. G. ESCRT-III controls nuclear envelope reformation. Nature 522, 236–239 (2015).

12. Jimenez, A. J. et al. ESCRT machinery is required for plasma membrane repair. Science 343, 1247136 (2014).

13. Skowyra, M. L., Schlesinger, P. H., Naismith, T. V & Hanson, P. I. Triggered recruitment of ESCRT machinery promotes endolysosomal repair. Science (1979) 360, eaar5078 (2018).

14. Radulovic, M. et al. ESCRT-mediated lysosome repair precedes lysophagy and promotes cell survival. EMBO J 37, e99753 (2018).

15. Schöneberg, J. et al. ATP-dependent force generation and membrane scission by ESCRT-III and Vps4. Science 362, 1423–1428 (2018).

16. Pfitzner, A.-K. et al. An ESCRT-III Polymerization Sequence Drives Membrane Deformation and Fission. Cell 182, 1140–1155.e18 (2020).

17. Schöneberg, J., Lee, I.-H., Iwasa, J. H. & Hurley, J. H. Reverse-topology membrane scission by the ESCRT proteins. Nat Rev Mol Cell Biol 18, 5–17 (2017).

18. McCullough, J. et al. Structure and membrane remodeling activity of ESCRT-III helical polymers. Science (1979) 350, 1548–1551 (2015).

19. Nguyen, H. C. et al. Membrane constriction and thinning by sequential ESCRT-III polymerization. Nat Struct Mol Biol 27, 392–399 (2020).

20. Talledge, N., et al. The ESCRT-III proteins IST1 and CHMP1B assemble around nucleic acids. bioRxiv 386532 (2018) doi:10.1101/386532.

21. Allison, R. et al. An ESCRT–spastin interaction promotes fission of recycling tubules from the endosome. Journal of Cell Biology 202, 527–543 (2013).

22. Allison, R. et al. Defects in ER–endosome contacts impact lysosome function in hereditary spastic paraplegia. Journal of Cell Biology 216, 1337–1355 (2017).

23. Sanjuan, M. A. et al. Toll-like receptor signalling in macrophages links the autophagy pathway to phagocytosis. Nature 450, 1253–1257 (2007).

24. Florey, O., Gammoh, N., Kim, S. E., Jiang, X. & Overholtzer, M. V-ATPase and osmotic imbalances activate endolysosomal LC3 lipidation. Autophagy 11, 88–99 (2015).

25. Martinez, J. et al. Molecular characterization of LC3-associated phagocytosis reveals distinct roles for Rubicon, NOX2 and autophagy proteins. Nat Cell Biol 17, 893–906 (2015).

26. Heckmann, B. L. et al. LC3-associated endocytosis facilitates β-amyloid clearance and mitigates neurodegeneration in murine Alzheimer’s disease. Cell 178, 536–551 (2019).

27. Nakamura, S. et al. LC3 lipidation is essential for TFEB activation during the lysosomal damage response to kidney injury. Nat Cell Biol 22, 1252–1263 (2020).

28. Corkery, D. P. et al. Vibrio cholerae cytotoxin MakA induces noncanonical autophagy resulting in the spatial inhibition of canonical autophagy. J Cell Sci 134, (2021).

29. Hooper, K. M. et al. V-ATPase is a universal regulator of LC3-associated phagocytosis and non-canonical autophagy. J Cell Biol 221, (2022).

30. Timimi, L. et al. The V-ATPase complex regulates non-canonical Atg8-family protein lipidation through ATG16L1 recruitment. Autophagy 18, 707–708 (2022).

31. Galluzzi, L. & Green, D. R. Autophagy-Independent Functions of the Autophagy Machinery. Cell 177, 1682–1699 (2019).

32. Lystad & Simonsen. Mechanisms and Pathophysiological Roles of the ATG8 Conjugation Machinery. Cells 8, 973 (2019).

33. Deretic, V. & Lazarou, M. A guide to membrane atg8ylation and autophagy with reflections on immunity. J Cell Biol 221, (2022).

34. Jia, X. et al. V. cholerae MakA is a cholesterol-binding pore-forming toxin that induces non-canonical autophagy. J Cell Biol 221, (2022).

35. Konstantinidis, G., Sievers, S. & Wu, Y.-W. Identification of Novel Autophagy Inhibitors via Cell-Based High-Content Screening. Methods Mol Biol 1854, 187–195 (2019).

36. Laraia, L. et al. The cholesterol transfer protein GRAMD1A regulates autophagosome biogenesis. Nat Chem Biol 15, 710–720 (2019).

37. Niggemeyer, G., et al. Synthesis of 20-Membered Macrocyclic Pseudo-Natural Products Yields Inducers of LC3 Lipidation. Angewandte Chemie International Edition 61, e202114328 (2022).

38. Gaetani, M. et al. Proteome integral solubility alteration: a high-throughput proteomics assay for target deconvolution. J Proteome Res 18, 4027–4037 (2019).

39. Xiao, J. et al. Structural basis of Ist1 function and Ist1-Did2 interaction in the multivesicular body pathway and cytokinesis. Mol Biol Cell 20, 3514–24 (2009).

40. Lata, S. et al. Helical structures of ESCRT-III are disassembled by VPS4. Science 321, 1354–7 (2008).

41. Bajorek, M. et al. Biochemical analyses of human IST1 and its function in cytokinesis. Mol Biol Cell 20, 1360–73 (2009).

42. Agromayor, M. et al. Essential role of hIST1 in cytokinesis. Mol Biol Cell 20, 1374–87 (2009).

43. Lee, S. et al. MITD1 is recruited to midbodies by ESCRT-III and participates in cytokinesis. Mol Biol Cell 23, 4347–4361 (2012).

44. Yang, D. et al. Structural basis for midbody targeting of spastin by the ESCRT-III protein CHMP1B. Nat Struct Mol Biol 15, 1278–86 (2008).

45. Babst, M., Katzmann, D. J., Estepa-Sabal, E. J., Meerloo, T. & Emr, S. D. Escrt-III: an endosome-associated heterooligomeric protein complex required for mvb sorting. Dev Cell 3, 271–82 (2002).

46. Katzmann, D. J., Babst, M. & Emr, S. D. Ubiquitin-dependent sorting into the multivesicular body pathway requires the function of a conserved endosomal protein sorting complex, ESCRT-I. Cell 106, 145–55 (2001).

47. van Niel, G., D’Angelo, G. & Raposo, G. Shedding light on the cell biology of extracellular vesicles. Nat Rev Mol Cell Biol 19, 213–228 (2018).

48. Laidlaw, K. M. E., Calder, G. & MacDonald, C. Recycling of cell surface membrane proteins from yeast endosomes is regulated by ubiquitinated Ist1. J Cell Biol 221, (2022).

49. Cada, A. K. et al. Friction-driven membrane scission by the human ESCRT-III proteins CHMP1B and IST1. Proc Natl Acad Sci USA 119, e2204536119 (2022).

50. Wong, S.-W., Sil, P. & Martinez, J. Rubicon: LC3-associated phagocytosis and beyond. FEBS J 285, 1379–1388 (2018).

51. Ulferts, R. et al. Subtractive CRISPR screen identifies the ATG16L1/vacuolar ATPase axis as required for non-canonical LC3 lipidation. Cell Rep 37, 109899 (2021).

52. Xu, Y. et al. ARF GTPases activate Salmonella effector SopF to ADP-ribosylate host V-ATPase and inhibit endomembrane damage-induced autophagy. Nat Struct Mol Biol 29, 67–77 (2022).

53. Babst, M., Wendland, B., Estepa, E. J. & Emr, S. D. The Vps4p AAA ATPase regulates membrane association of a Vps protein complex required for normal endosome function. EMBO J 17, 2982–93 (1998).

54. Chang, C.-L. et al. Spastin tethers lipid droplets to peroxisomes and directs fatty acid trafficking through ESCRT-III. J Cell Biol 218, 2583–2599 (2019).

55. Mast, F. D. et al. ESCRT-III is required for scissioning new peroxisomes from the endoplasmic reticulum. J Cell Biol 217, 2087–2102 (2018).

56. Khan, I. & Steeg, P. S. Endocytosis: a pivotal pathway for regulating metastasis. Br J Cancer 124, 66–75 (2021).

57. van Weert, A. W., Geuze, H. J., Groothuis, B. & Stoorvogel, W. Primaquine interferes with membrane recycling from endosomes to the plasma membrane through a direct interaction with endosomes which does not involve neutralisation of endosomal pH nor osmotic swelling of endosomes. Eur J Cell Biol 79, 394–9 (2000).

58. Kim, J.-H., Choi, H.-S. & Lee, D.-S. Primaquine Inhibits the Endosomal Trafficking and Nuclear Localization of EGFR and Induces the Apoptosis of Breast Cancer Cells by Nuclear EGFR/Stat3-Mediated c-Myc Downregulation. Int J Mol Sci 22, (2021).

59. Mishra, A., Hourigan, D. & Lindsay, A. J. Inhibition of the endosomal recycling pathway downregulates HER2 activation and overcomes resistance to tyrosine kinase inhibitors in HER2-positive breast cancer. Cancer Lett 529, 153–167 (2022).

60. Peña-Martinez, C., Rickman, A. D. & Heckmann, B. L. Beyond autophagy: LC3-associated phagocytosis and endocytosis. Sci Adv 8, eabn1702 (2022).

61. Gaetani, M. & Zubarev, R. A. Proteome Integral Solubility Alteration (PISA) for High-Throughput Ligand Target Deconvolution with Increased Statistical Significance and Reduced Sample Amount. Methods Mol Biol 2554, 91–106 (2023).

62. Zhang, X., Lytovchenko, O., Lundström, S. L., Zubarev, R. A. & Gaetani, M. Proteome Integral Solubility Alteration (PISA) Assay in Mammalian Cells for Deep, High-Confidence, and High-Throughput Target Deconvolution. Bio Protoc 12, (2022).

63. Cox, J. & Mann, M. MaxQuant enables high peptide identification rates, individualized p.p.b.-range mass accuracies and proteome-wide protein quantification. Nat Biotechnol 26, 1367–72 (2008).

64. Xu, Y. et al. A Bacterial Effector Reveals the V-ATPase-ATG16L1 Axis that Initiates Xenophagy. Cell 178, 552–566.e20 (2019).

65. Nakamura, S. et al. LC3 lipidation is essential for TFEB activation during the lysosomal damage response to kidney injury. Nat Cell Biol 22, 1252–1263 (2020).

